# Linking offline learning mechanisms to anxiety traits

**DOI:** 10.1101/2023.06.21.546031

**Authors:** Qianqian Yu, Yuejia Luo, Ray Dolan, Jianxin Ou, Chuwen Huang, Haiteng Wang, Zhibing Xiao, Matthew Nour, Yunzhe Liu

**Author notes:** Correspondence (Y.L.). These authors contributed equally (Qianqian Yu, Yuejia Luo).

## Abstract

Understanding how we learn about the value and structure of our environment is central to neurocognitive theories of many psychiatric and neurological disorders. Learning processes have been extensively studied during performance of behavioural tasks (online learning) but less so in relation to “resting” (offline) states. A candidate mechanism for such offline learning is “replay,” the sequential neural reactivation of past experiences. Notably, value-based learning is especially tied to replay unfolding in reverse order relative to the original experience (“backward replay”). Here, we demonstrate the utility of EEG-based neural decoding for investigating offline learning, and relate it to trait anxiety. Participants were first required to infer sequential relationships among task objects by using a learned rule to reorganise their visual experiences into distinct sequences. Afterwards, they observed that the final object in one of the sequences was associated with a monetary reward and then entered a post-value resting state. During this rest, we found evidence of backward replay for reward-linked object sequences. The strength of such replay was negatively associated with trait anxiety and positively predicted an increased behavioural preference for reward-predictive stimuli. We also found that high-trait-anxiety individuals showed inefficient credit assignment irrespective of reward magnitude, indicating that this effect does not merely reflect reduced reward sensitivity. Together, these findings suggest a potential aberrant replay mechanism during offline learning in individuals with high trait anxiety. More broadly, our approach illustrates the potential of EEG for measuring structured neural representations in vivo, with implications for studying cognition across a range of neuropsychiatric and neurological disorders.

## Introduction

Mechanistic accounts of psychopathological symptoms often centre on maladaptive learning processes, such as structure learning – understanding how entities are related^1^, and value learning – assigning value to states or actions^2,3^. For instance, anxiety has been linked to a bias prioritizing punishment over reward signal, which negatively impacts value learning^4,5^, and may lead to avoidant behaviours that limit engagement in potentially rewarding activities^6–11^. While most research has focused on online (i.e., on-task) learning, recent work emphasises the importance of offline learning that occurs during rest and sleep^12^.

Offline learning, as occurs during rest or sleep, involves processing and consolidation of information acquired during an awake state^13,14^. A key neural mechanism of offline learning is neural “replay,” the spontaneous sequential reactivation of on-task neural representations^15–17^. Replay is crucial for both structure learning, where internal relationships are encoded^1,18^, and value learning^19,20^, in which reward signals are assigned to relevant states^21–23^. Of particular interest is backward replay – reactivating experiences in reverse order – which is highly responsive to reward^19,24^, and enables non-local value learning^25,26^, allowing reward signals to propagate to preceding (non-local) states in a sequence. Disruptions in such offline replay processes may underlie dysfunctional reward processing in psychopathologies^27^.

Recent methodological advances enable the non-invasive detection of replay-like sequential reactivations in humans^28^, typically through magnetoencephalography (MEG) ^1,18,29–32^ or functional magnetic resonance imaging (fMRI) ^33,34^. Studies using these techniques indicate that reward can enhance backward replay of learned sequences^18,22,29^, suggesting that replay is integral to the propagation of reward signals beyond direct reward associations.

Trait anxiety is increasingly viewed as a biomarker or vulnerable phenotype for mood disorders, encompassing both depression and anxiety^35,36^. Rather than separating anxiety from depression, we examined whether offline replay mechanisms were disrupted in individuals with higher trait anxiety, particularly in the context of non-local value learning. We focused on reward-based credit assignment, for which backward replay serves as a robust neural signature^18,19,24^. Building on our previous MEG study^1,18^, we employed an established sequence-learning paradigm to investigate these processes in individuals with varying levels of trait anxiety, but now using EEG.

Participants first learned the structure of two picture sequences and then underwent a value-learning phase where the final image of one sequence was paired with a monetary reward, while the other sequence had no such pairing. Preference ratings collected before and after this phase served as an implicit behavioural measure of credit assignment. Crucially, prior to value learning, we ensured all participants had learned the “true” sequence structure to a similar extent, so that any observed deficits in reward-based preference shifts and replay mechanisms could not be attributed to poor structural knowledge.

We recorded electroencephalography (EEG) – a method more widely accessible than MEG in clinical settings – to detect replay during rest following value learning. All 80 subjects were carefully recruited to ensure an even sampling across levels of trait anxiety, as assessed by the Chinese-validated version of Spielberger Trait Anxiety Inventory (STAI) ^37,38^. We hypothesised that higher trait anxiety would be associated with weaker backward replay of reward-linked sequences, accompanied by smaller preference shifts reflecting impaired value updating.

To rule out the possibility that high trait anxiety simply reflects diminished reward sensitivity, we conducted a separate behavioural experiment (n = 200) in which we manipulated reward magnitude (¥10 vs. ¥100). Across both reward conditions, individuals with higher trait anxiety exhibited diminished propagation of reward value through the learned sequence. Together, this outcome suggests that aberrant offline replay mechanisms may underlie value learning deficits beyond mere reward sensitivity, providing insights into the maladaptive learning patterns associated with trait anxiety.

## Results

### Task

Participants completed the task over two study days. On Day 1 (behavioural training only), they learned only the task structure. Eight images were presented in two scrambled visual sequences, [B’ A D’ B] and [A’ C C’ D], while the true latent orders were A → B → C → D and A’ → B’ → C’ → D’. An explicit “unscrambling” rule, for example “the first picture of visual sequence 1 is actually the second picture of structural sequence 2”, taught participants to map the visual order onto the hidden structural order. Crucially, the structural neighbours that later define replay transitions, such as A → B, never appeared consecutively in the visual streams, so any A → B reactivation detected at rest must reflect internally inferred structure rather than replay of a perceptual pairing. Following our previous MEG work^1,18^, this procedure was designed to induce off-task replay of inferred transitions between stimuli (i.e., beyond simple experience), as well as to track how reward information propagates along these inferred sequences. This study focused on distinguishing replay of reward-associated versus neutral sequences. Because replay and spontaneous memory reactivation can also prioritise weakly learnt experiences^39–41^, participants were required to reach ≥80% accuracy on the Day 1 structure learning task to minimise confounds from incomplete structural knowledge. Only those meeting this threshold proceeded to the EEG experiment on Day 2.

On Day 2, the task involved a sequence learning phase with a new set of eight visual stimuli, mirroring the structure learning task from Day 1. This session was conducted during a 64-channel EEG scan. Initially, participants rated their liking for each stimulus on a 1-9 scale, followed by a functional localiser task wherein each picture was displayed randomly (crucial for neural state decoding). Following the functional localiser, participants completed a sequence learning phase where they learned how 8 new pictures related to 2 underlying structural sequences, using the same rule as learnt on Day 1. Participants then entered a value learning phase, where the final picture in one of the structural sequences (D) was associated with reward, while the other (D’) was associated with a neutral state. The task concluded with a “position probe” session, where participants identified each picture’s position in the sequence, followed by a final preference rating. A preference change was used to quantify the impact of reward on subjective valuation^42^. Three resting state sessions (4 minutes each) were included: one at the task’s start, prior to structure learning (pre-task); and surrounding the value learning phase (pre-value and post-value), to capture spontaneous replay of the task sequences. For further details, refer to the Methods section.

### Behavioural performance

Seven participants were excluded for failing Day 1 structure learning, and an additional five were excluded due to excessive movements and/or EEG artifacts, leaving 68 participants in the final sample. Behavioural analysis confirmed these 68 participants successfully learned the structural task sequences. The overall mean trait anxiety score was 42.57 ± 1.34. Furthermore, as expected, our trait anxiety measure correlated with other anxiety questionnaires (e.g., the Self-Rating Anxiety Scale^43^ (SAS; *r* = 0.65, *p* < 0.001), Penn State Worry Questionnaire^44,45^ (PSWQ; *r* = 0.65, *p* < 0.001), and Intolerance of Uncertainty Scale^46,47^ (IUS; *r* = 0.37, *p* = 0.002)), demonstrating construct validity. Trait anxiety also correlated with depression scores, such as the Self-Rating Depression Scale^48^ (SDS; *r* = 0.81, *p* < 0.001), in line with the high comorbidity between trait anxiety and depression well-documented in epidemiological studies^49,50^.

The main EEG task consisted of four phases (Figure 1). During the functional localiser phase, sequence pictures were presented randomly to train neural decoders. To ensure attention, stimuli were occasionally (20% probability) shown upside down, requiring a button response from subjects. The mean response accuracy for this task was 97.54 ± 0.21%.

**Fig. 1.**
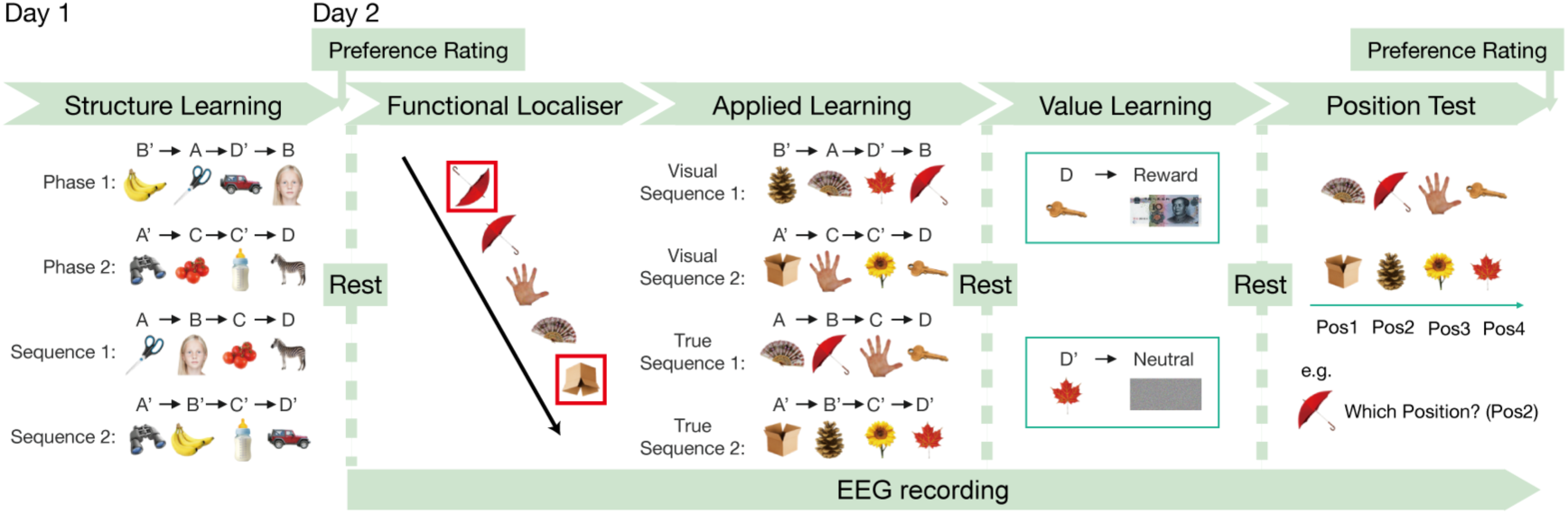
Task Design. On Day 1, during a structure learning phase, participants were trained on the mapping rule between the order of stimuli presentation (the “visual sequence”, e.g., “B’->A->D’->B”) and the task-relevant order (the “true sequence”, e.g., “A->B->C->D”) (details not shown). On Day 2, they were exposed to a new set of stimuli but followed the same mapping rule. Stimuli preferences were rated twice: once at the beginning and again at the end of the task. Subjects engaged in the sequence learning task ^18^ while undergoing whole-brain EEG recording. During the functional localiser phase, stimuli were presented in random order to train decoders. In the sequence learning phase, stimuli were presented in the same manner as on Day 1, with subjects required to apply the learned structure to reorder the new stimuli. Following this phase, the last stimulus of each “true sequence”, either D or D’, was paired with a reward or neutral icon. Finally, subjects were probed on the position of all stimuli in the true sequence. There were three resting states throughout the task, used to capture spontaneous replay of the task sequences. During these resting states, subjects were instructed to keep their eyes open for 4 minutes without engaging in any specific task.

Next, in the sequence learning phase, participants applied the Day 1 mapping rule to newly introduced Day 2 stimuli. After each learning run, their knowledge of the true stimulus order was assessed without feedback, revealing a mean accuracy of 84.58 ± 1.47% (chance level = 50%). Accuracy improved over the course of the learning runs (*F* (2, 134) = 58.59, *p* < 0.001, *η²* = 0.47). Notably, no correlations emerged between trait anxiety scores and any learning or test performance measures (*r*s < 0.10, *p*s > 0.20), suggesting that high-vs. low-trait-anxiety individuals did not differ in acquiring task contingencies (i.e., applying the abstracted structure to new sensory inputs). Following Nour et al. (2021), we quantified learning efficiency by calculating a “learning lag,” which regresses quiz performance on run number, where larger values indicate slower performance improvements (i.e., lower efficiency). No significant correlations were observed between learning lag and trait anxiety (accuracy: *r* = −0.06, *p* = 0.586; response time: *r* =-0.10, *p* = 0.431), indicating that trait anxiety did not impact the transfer of task-structure knowledge to new experiences in this study.

In the value learning phase, the last object of each true sequence (D or D’) was paired with a reward or a neutral icon, respectively. Participants pressed specific buttons to indicate their recognition of reward or non-reward, achieving an overall mean accuracy of 94.42 ± 1.91%. The position probe then tested subjects on the ordinal position of each picture within the learned sequence, revealing an overall mean accuracy of 96.98 ± 0.86%, indicating correct encoding of the task structure. See Methods section for details of behavioural testing procedures.

At the end of the entire task, participants again rated their preference for stimuli. ANOVA analysis, considering factors of structural position, time (pre-vs. post-task), and sequence (reward vs. neutral), revealed a 3-way interaction (position x time x sequence) on final preference ratings (*F* (3, 201) = 4.82, *p* = 0.003, *η*² = 0.067), such that maximal preference was for the final picture of the rewarded sequence, after value learning. Specifically, for the rewarded sequence, an interaction between position and time was observed (*F* (3, 201) = 5.44, *p* = 0.001, *η*² = 0.075, Fig. 2a), with a notable preference increase for the last object post-task (pre: 5.63 ± 0.21, post: 6.03 ± 0.23, *t* (67) =-3.32, *p* = 0.001, Fig. 2a). No such effects were found for the neutral sequence. Additionally, a significant interaction between sequence and time was noted (*F* (1, 67) = 4.93, *p* = 0.03, *η*² = 0.068), with a marginally higher post-task preference rating for the rewarded sequence compared to the neutral sequence (*p* = 0.079).

**Fig. 2.**
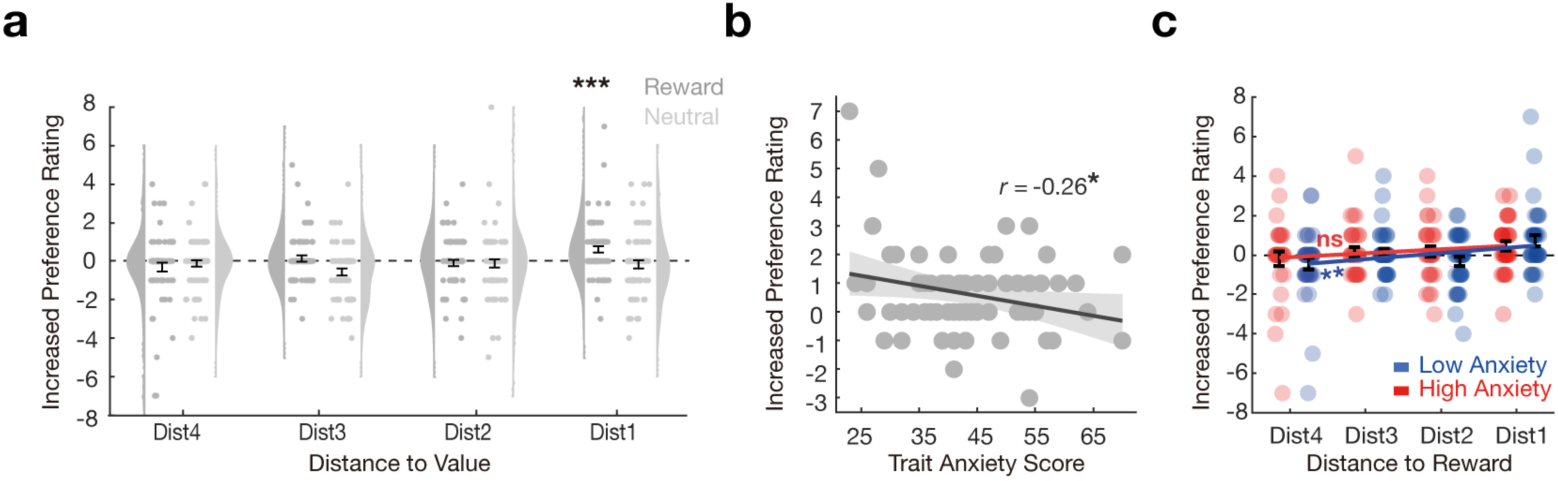
Change of Preference Rating following Value Learning. a, Increased (post - pre) preference rating of stimuli based on distance to reward/neutral icon (referred to as sequence position) for both reward and neutral sequences. **b,** The increased preference rating on object (D), paired with reward, is linked to trait anxiety score evident in a negative correlation between increased preference rating and trait anxiety score. **c,** Increases in stimuli preference as a function of distance to reward. There was a significant positive linear relationship between closeness to reward and increase in preference for low anxious subjects, but not so for subjects with high anxiety (no significant difference between groups). Error bars show SEM; each dot indicates results from one subject. The solid line reflects the best robust linear fit, \**p* < 0.05, \*\**p* < 0.01, \*\*\**p* < 0.001.

Importantly, we found a significant negative correlation between trait anxiety score and increased (post - pre) preference rating for the rewarded stimulus (*r* =-0.26, *p* = 0.036, Fig. 2b), indicating a blunting of a preference increase in the more anxious individuals. This finding aligns with previous research linking trait anxiety to impaired value learning^4,5^. Furthermore, when treating anxiety as a categorical variable (below vs above the clinical severity cut-off, based on prior work^38,51^, we observed that for low anxious individuals, the preference for all reward-predictive stimuli increased proportionally to their proximity to the reward (*β* = 0.31, *p* = 0.008, Fig. 2c), an effect not evident in high anxious individuals (*β* = 0.20, *p* = 0.106), although the difference between groups was not significant. This suggests that trait anxiety may be associated with impairments in value updating following experience, a process likely to rely on an internal model of the task, namely the actual reward sequence.

### Neural decoding

Having uncovered a significant relationship between post-value learning behavioural preference (a reflection of value learning) and anxiety, we next investigated potential neural correlates of learning. A post-value rest period separated value-learning and preference ratings, allowing us to specifically investigate post-learning offline neural replay.

To measure sequential replay during rest, we first trained multivariate neural decoders for each task picture, based on visually evoked response EEG data from the functional localiser, using lasso-regularised logistic regression trained in k-fold cross-validation. Importantly, as the localiser was conducted at the start of each scanning session, prior to any learning, this removed any possibility that information pertaining to task-structure might be present in the classifiers. Consistent with prior studies^1,18^, we trained a distinct set of decoders from neural data at each 10 ms window following stimulus onset. Classification accuracy in held-out data was evaluated using a 10-fold cross-validation^52^. The peak decoding accuracy was observed at 180 ms post-stimulus onset (23.21 ± 0.90%, one-sample *t*-test, *t* (67) = 11.934, *p* < 0.001; chance level: 12.50%, Fig. 2a), as in our prior work^1^. Importantly, there was no correlation between the peak decoding accuracy and trait anxiety scores (*r* = 0.08, *p =* 0.50), which might otherwise confound later replay-anxiety relationships. The classifiers demonstrated high specificity for task states, as the predicted probability of the trained models significantly exceeded the threshold only when the test state matched the state the model was trained to detect (Supplementary Fig. 1). For all subsequent analyses, we used stimulus classifiers trained at 180 ms post-stimulus onset.

### Selective increase of backward replay after value learning

We assessed evidence for spontaneous neural replay of inferred sequences in resting-state EEG data using Temporal Delayed Linear Modelling (TDLM)^28^, consistent with methodologies used in prior MEG studies^1,18,29,31^. TDLM captures unique neural patterns while accounting for co-activations and autocorrelations among all states, thereby minimising false positives arising from overlapping image representations^28^. This involves applying trained classifiers to the neural (time, sensors) matrix to generate a (time, states) reactivation probability matrix. We then quantified the degree to which these reactivation probabilities systematically adhered to a specified task transition matrix at various “replay lags” (speeds, 10 ms – 600 ms) using generalised linear models, for each participant separately. We measured “replay strength” across two stimuli orders: forward (e.g., [A->B]) and backward (e.g., [B->A])^18,29^. Notably, previous studies using MEG have revealed an increase in backward replay following receipt of rewards^18,22,29^.

We found significant backward replay for the rewarded sequence in the rest period following value learning (post-value), with a state-state lag (speed) of 50-80 ms time lags, peaking at 60 ms (*β* = 0.02 ± 0.01, one-sample *t*-test, *t* (67) = 3.20, *p =* 0.001, significance threshold from a non-parametric permutation test, FWE corrected across all lags, Fig. 3b). Importantly, no such backward replay was observed in the reward sequence during the rest period before value learning (pre-value) or during the rest period at the start of the task (pre-task). Conversely, we found no significant backward replay for the neutral sequence in any of the three rest periods (Supplementary Fig. 2).

**Fig. 3.**
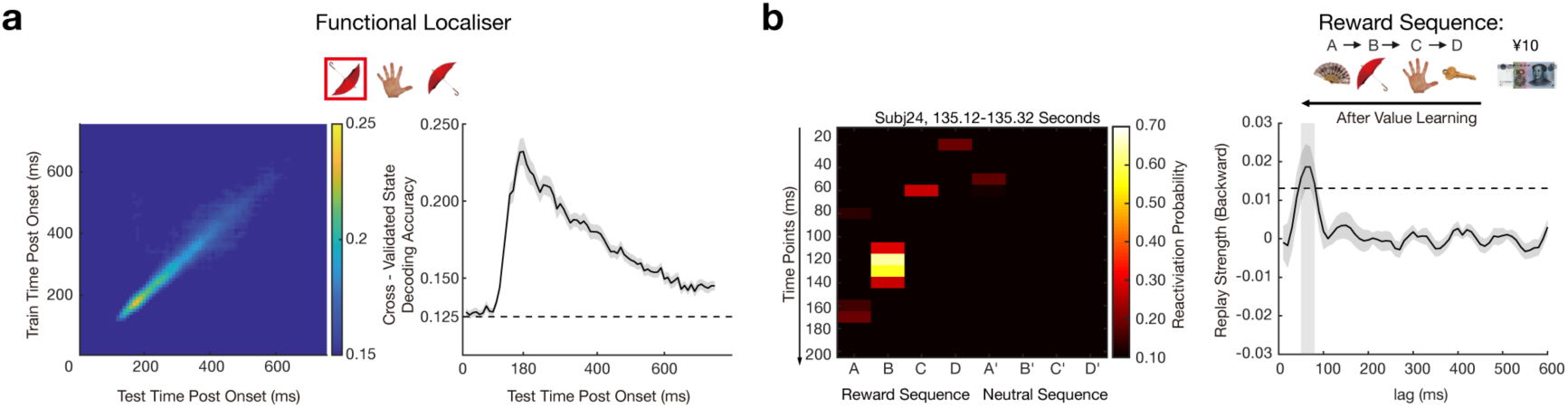
Neural Decoding and Backward Replay After Value Learning. **a**, Cross-validated decoding accuracy of 8 stimuli based on whole-brain EEG signal in the functional localiser phase. As with Liu et al.^18^, a binary classifier was trained independently for each stimulus using logistic regression. On the left, it is the decoding accuracy of classifiers trained on one time point (row) and tested on all time points (column) from stimuli onset to post 750 ms. On the right, the decoding time course of classifiers trained and tested at the same time. The horizontal dashed line represents the peak-level *P_FWE_*= 0.05 threshold for chance-level accuracy (test-data label permutation). Decoding performance for each state is shown in Supplementary Fig. 1. **b,** An example of replay events from an exemplar subject is shown for visualization purposes alone (left panel). Each row depicts reactivation probabilities of all stimuli at a given time point. On the right panel is the mean backward replay for the reward sequence, after value learning. The grey shaded area indicates significant time lags of replay. This result is similar with previous studies measuring replay in MEG^18,29^.

We found a significant interaction between time (pre vs. post-value learning) and sequence type (rewarded vs. neutral) in the magnitude of backward replay (measured at the maximum time lag (70 ms) averaging across both reward and neutral sequences: *F* (1,67) = 6.80, *p* = 0.011, *η*² = 0.092, Reward: β*_pre_* = 0.006 ± 0.0042, β*_post_* = 0.0187 ± 0.0053, paired *t*-test, *t* (67) =-3.131, *p* = 0.003, Neutral: β*_pre_* = 0.0028 ± 0.0044, β*_post_* =-0.0036 ± 0.0055, paired *t*-test, *t* (67) = 1.301, *p* = 0.198). This indicates a significant increase in backward replay after value learning for the rewarded compared to the neutral sequence. As anticipated, we found no replay of the visual sequence (observed picture transitions, not part of the latent structural sequence) in any rest session (Supplementary Fig. 3). In contrast to prior work^1,18^, we did not observe forward replay in either the rewarded or neutral sequences during any rest period that exceeded *P_FWE_* < 0.05 threshold (Supplementary Fig. 4). See Supplementary Fig. 5-6 for additional validation results. We also trained classifiers on post-learning data (during the position task), which yielded a lower decoding accuracy (21.40 ± 0.96%) than the pre-task localiser-trained classifiers (23.18 ± 0.86%; paired *t-*test, *t* (67) = −1.79, *p* = 0.039). No replay was observed in this condition, likely because task-related representational overlap reduced decoding accuracy and resulted in non-significant replay measures.

To test whether learning produced a selective blurring of neighbouring states that might masquerade as replay—e.g., if the pattern for state A became more like state B and thus yielded an apparent A → B replay during rest—we carried out two checks. First, a cross-temporal decoding analysis showed that classifiers trained on pre-learning functional-localiser data at 180 ms remained highly discriminative when applied to signals from the final position-test phase (Supplementary Fig. 7a). This stable, state-specific generalisation argues against widespread representational drift after learning. Second, we asked whether decoders misclassified adjacent items more often than non-adjacent ones (for example, whether a classifier for state A was more likely to predict state B than state C). In the position test, we modeled the decoder probabilty for each non-target state (i.e., confusion), using a linear mixed-effects model^53–55^, Probability Confusion ∼ Adjacency ∗ Time + (Adjacency ∗ Time | Subject). Adjacency was not significant for the reward sequence (*β* =-0.0008 ± 0.003, *p* = 0.911) or the neutral sequence (*β* = 0.0003 ± 0.004, *p* = 0.942), and there were no adjacency-by-time interactions (both *p* > 0.70; Supplementary Fig. 7b), indicating neighbouring items did not become disproportionately confusable after learning, such that representational drift cannot account for the selective backward replay observed for the reward sequence.

### High trait anxiety is associated with reduced backward replay

We observed a significant negative correlation between trait anxiety and an increase in backward replay for the rewarded sequence after value learning (*r* =-0.27, *p* = 0.029, Fig. 4a left panel, also shown for low vs. high anxiety comparison on the right panel). Moreover, this increase in backward replay was negatively correlated with SDS scores (*r* = −0.29, *p* = 0.015), and marginally correlated with SAS scores (*r* = −0.220, *p* = 0.07) and PSWQ scores (*r* = −0.22, *p* = 0.066). These findings suggest that deficits in offline value learning may be broadly linked to mood-related pathology.

**Fig. 4.**
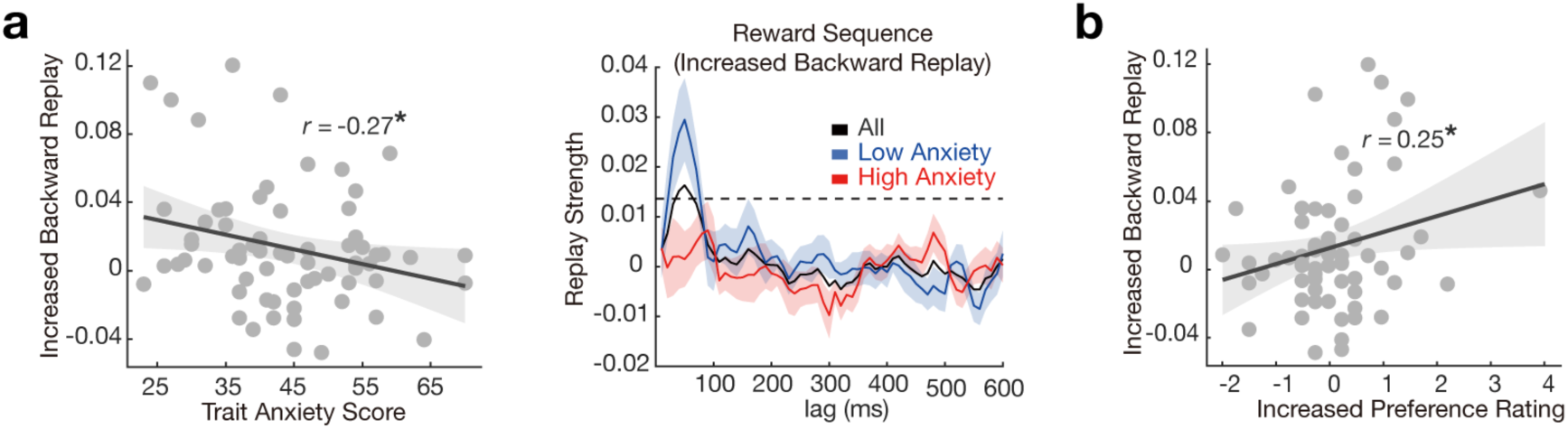
Diminished Reward Effect on Backward Replay is Associated with Trait Anxiety. a, A selective increase in backward replay after value learning is negatively correlated with individual trait anxiety. The scatter plot is shown on the left, and the low vs. high anxiety group comparison is shown on the right. The dashed line is the *P_FWE_*= 0.05 permutation threshold after controlling for multiple comparisons. **b,** The reward effect on replay is linked to behavioural change - there is a positive correlation between increase in backward replay and an increase in preference rating, specifically for reward sequence. Each dot is a subject, the solid line represents the best linear fit with shaded area indicating SEM. \**p* < 0.05.

We also found a positive correlation between the increase in backward replay for the reward sequence and changes in preference ratings from pre-to post-task (*r* = 0.25, *p* = 0.042, Fig. 4b). Additionally, in the reward sequence, an interaction effect was observed between group (high vs. low trait anxiety) and time (pre vs. post value learning) for (increase in) backward replay (*F* (1, 66) = 4.77, *p* = 0.032, *η*² = 0.067). Specifically, individuals with low trait anxiety demonstrated a larger increase in backward replay following value learning than did those with high trait anxiety (β_*$+_ = 0.02 ± 0.01, β_-./-_ = 0.00 ± 0.01, two-sample *t*-test, *t* (66) =-2.26, *p* = 0.013, Supplementary Fig. 8).

In contrast, we found neither significant correlations (*r*s > 0.1, *p*s >0.1) nor interaction effects (*F* (1, 66) = 1.69, *p* = 0.199, *η*² = 0.025) for backward replay changes in the neutral sequence (pre-vs. post-value learning) and trait anxiety or scores on other mood-related questionnaires.

One might argue that a neutral outcome in our study could be perceived as relatively aversive. Individuals with higher trait anxiety are often thought to “ruminate” on negative outcomes^56,57^, and if such rumination is expressed via replay during rest^27^, one might predict a positive correlation between neutral-sequence replay strength and anxiety traits. In a post-hoc analysis, we observed a marginally significant positive correlation between post-value-learning backward replay of neutral sequences (averaged over the 20–60 ms time lag, following previous human replay studies^1,18^) and trait anxiety (*r* = 0.24, *p* = 0.051). Taken together, these results indicate that high trait anxiety specifically impairs backward replay in reward-associated sequences. More speculatively, individuals with higher trait anxiety may engage in a form of “rumination” by replaying more of neutral sequences as if they were less favourable experiences.

### Reward-based neural map representation is positively related to backward replay, negatively linked to trait anxiety

Finally, we explored whether diminished reward-associated backward replay in subjects with high trait anxiety was linked to a distorted neural representation of task structure. For this, we implemented a representational similarity analysis (RSA), similar to approaches used in previous studies^1,58^. The RSA quantifies the representational geometry of evoked neural responses both before and after value learning (utilizing data from the functional localiser and position probe sessions, respectively – see Methods and Figure 1). By contrasting representational patterns (representational dissimilarity matrices, RDMs) from before (functional localiser) and after (position probe) value learning, we generated a [state x state] “similarity change” matrix (RDM _(FL)_ - RDM _(pp),_ which is equivalent to RSM _(pp)_ – RSM _(FL)_, where RSM denotes representational similarity matrices) for each time point following stimulus onset.

These “similarity change” matrices were then regressed onto a task design matrix that included predictors for an abstracted “position” representation (1, 2, 3, 4) and two “sequence” representations (reward and neutral sequences, as illustrated in Fig. 5a, with additional details provided in the Methods section). The position matrix assigned a value of 1 to image pairs sharing the same ordinal position, and 0 otherwise, while each sequence matrix assigned 1 to image pairs within that same sequence. This allowed us to determine whether learning-related changes in similarity emerged because two images occupied the same position in a sequence, or because they both belonged to a reward or neutral sequence. Ultimately, this analysis enabled us to quantify the changes in task-specific neural representation induced by the learning process.

**Fig. 5.**
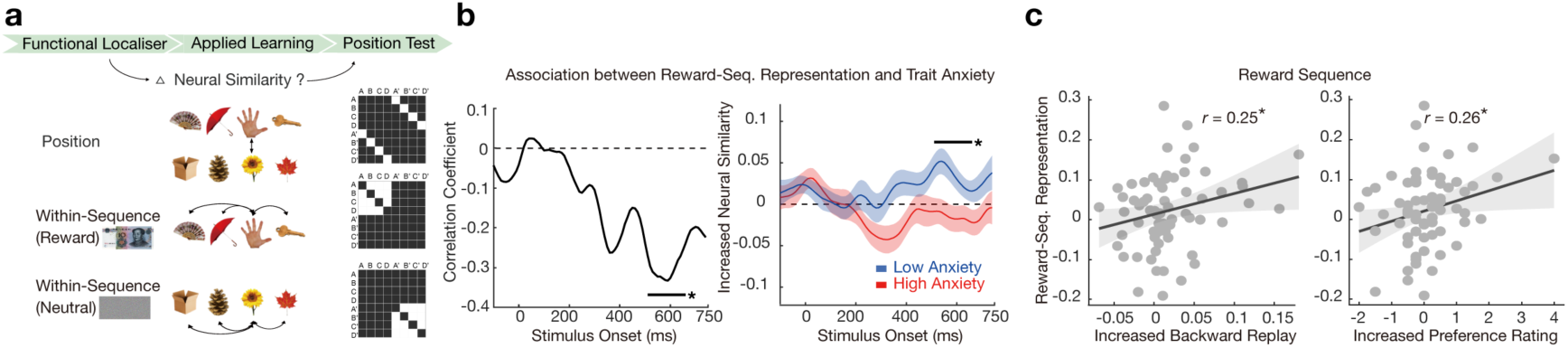
Neural Representation of Reward Sequence and Trait Anxiety. a, Hypothesised patterns of learning-induced “similarity change” and corresponding state x state design matrices are presented. **b,** The learning-induced change in the neural representation of the reward sequence is negatively correlated with trait anxiety scores (left panel). Notably, this change is observed in subjects with low anxiety alone but not in those with high anxiety (right panel). The group difference is also significant. The horizontal black line denotes the significant time window (cluster-level *P_FWE_* < 0.05) where the correlation between the reward representation and trait anxiety scores is significant. **c,** The neural representation of the reward sequence positively correlates with an increase in backward replay (at a 50 ms lag, the timepoint showing peak increase in replay from pre-to post-value rest) and with preference ratings, specifically for the reward sequence. Each dot represents a subject, with the solid line indicating the best linear fit and the shaded area representing the standard error of the mean (SEM). The shaded area depicts the standard error of the mean across subjects. \**p* < 0.05.

Given our focus on value learning, our primary hypotheses related to the reward representation regressor. In the context of our RSA analysis, we operationalised a reward representation as a neural index which quantifies the degree to which the neural patterns for states within the rewarded sequence become more similar from the start to the end of the task. We found no significant evidence for the emergence of reward representation across all participants following learning at any timepoint. However, in view of the above relationships between replay and anxiety, we also correlated the strength of reward representation at each time point post-stimulus onset with anxiety scores. A negative correlation emerged between reward representation and trait anxiety during 520-670 ms post-stimulus onset (cluster-level *P*_FWE_ = 0.05, non-parametric permutation test; peak 600 ms: *r* =-0.33, *p* = 0.0056, Fig. 5b left panel). Subsequent analysis of the mean reward representation in high and low anxiety groups (defined as above) revealed that only subjects with low trait anxiety showed a significant reward representation from 480-630 ms post-stimulus onset (cluster-level *P*_FWE_ = 0.02, non-parametric permutation test; peak 540 ms: *β* = 0.05 ± 0.02, one-sample *t*-test, *t* (35) = 3.37, *p* = 0.0009, Fig. 5b right panel). This effect was not significant in the high anxiety group (*β* =-0.01 ± 0.02). There was also a significant group difference between high vs. low anxiety subjects (two-sample *t*-test, *t* (66) = 2.77, *p* = 0.007). Furthermore, the magnitude of reward-related representational similarity positively correlated with the strength of reward-induced backward replay (*r* = 0.25, *p* = 0.038, Fig. 5c left panel), and also with an increase in preference rating for stimuli within the reward sequence (*r* = 0.26, *p* = 0.034, Fig. 5c right panel). Such effect was absent in relation to the neural representation of neutral sequences or representation of structural position (see Supplementary Fig. 9).

Finally, we evaluated the generalisability and potential clinical relevance of these findings by conducting a supplementary regression analysis that integrated behavioural markers (increased preference ratings for reward-predictive stimuli) and neural measures (representational change and increased backward replay strength in the reward sequence) to predict trait anxiety scores. Using a general linear model (GLM), with leave-one-out cross-validation, we observed a modest but significant correlation between predicted and actual anxiety scores (*r* = 0.32, *p* = 0.009). By contrast, a model using only behavioural predictors failed to reach significance (*r* = 0.13, *p* = 0.291), whereas a model using only neural predictors showed a significant correlation (*r* = 0.31, *p* = 0.009). Furthermore, in a combined model, both increased backward replay (*β* = – 2.95, *p* = 0.025) and representational change of the reward sequence (*β* = –2.66, *p* = 0.042) emerged as significant predictors. These results suggest that neural markers substantially enhance the prediction of trait anxiety, implying that either a combination of behavioural and neural measures or neural measures alone can explain a meaningful portion of variance in trait anxiety. The accessibility and cost-effectiveness of scalp EEG underscore the feasibility of applying these biomarkers in diverse research and clinical contexts.

### Reward sensitivity and trait anxiety on preference change

To rule out the possibility that the effects in our EEG study could be attributed to reduced reward sensitivity, we conducted a separate behavioural experiment (n = 200) manipulating reward magnitude (¥10 vs. ¥100; Fig. 6a). Thirty-one participants were excluded for either failing to learn the task structure or being unable to complete the task, leaving a final sample of 169 subjects (106 females; mean age = 19.89 ± 2.03 years). Participants were again assessed by the Chinese-validated version of STAI^37,38,59,60^, with an overall mean trait anxiety score of 44.68 ± 8.78. The results showed that individuals with high trait anxiety exhibited inefficient credit assignment irrespective of the reward magnitude, suggesting that anxiety impairs value updating in an offline context rather than merely reflecting reduced reward sensitivity.

**Fig. 6.**
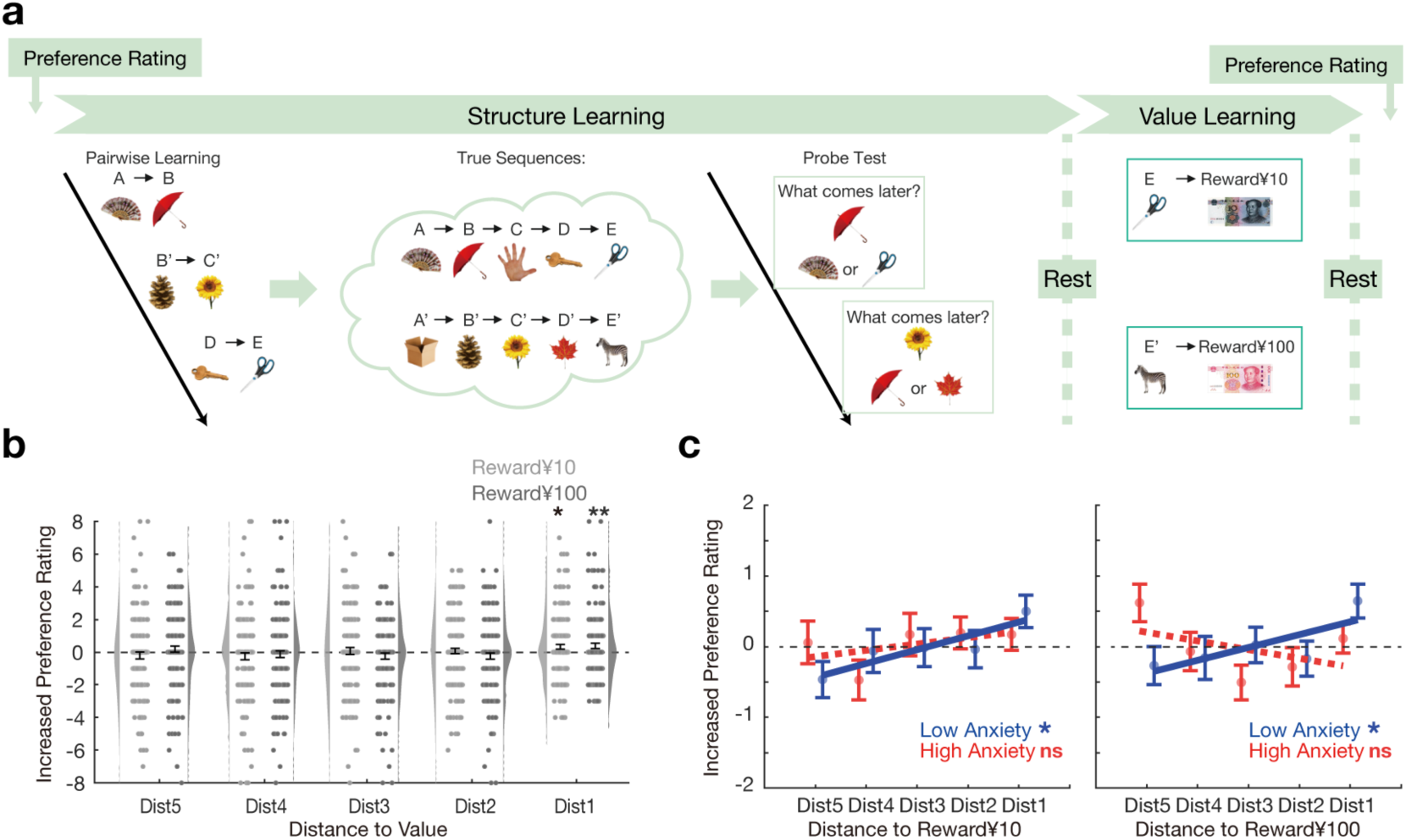
Change in Preference Ratings following Value Learning with Different Reward Magnitudes. a, Task design. Similar to the design of our EEG study, this behavioural experiment manipulated reward magnitude (¥10 vs. ¥100) to examine its effect on participants with varying levels of trait anxiety. Preferences were rated twice (pre-and post-task). The sequence learning phase involved pairwise training (e.g., B’→C’, A→B, D’→E’) and learning a “true sequence” (e.g., A→B→C→D→E). The final stimulus in each sequence (E or E’) was paired with either ¥10 or ¥100. Two resting states (4 minutes each) were included, mirroring the EEG study design. **b,** Mean increase (post minus pre) in preference ratings relative to distance from the reward icon (¥10 or ¥100). **c,** Stimulus preference increases as a function of proximity to the reward icon, separated by trait anxiety group (low vs. high) for ¥10 (left) and ¥100 (right). Only low-anxiety participants showed a significant positive linear relationship between closeness to reward and preference increase. Error bars show SEM; each dot indicates results from one subject. The solid line reflects the best robust linear fit, \**p* < 0.05, \*\**p* < 0.01.

Similar to the EEG experiment, participants performed both structure learning and value learning tasks (Fig. 6a). After rating their preferences for ten stimuli on a 1–9 scale (“strongly dislike” to “strongly like”) at the start of the experiment, they then underwent a pairwise learning session to infer two “true sequences” ([A/B/C/D/E] and [A’/B’/C’/D’/E’]). Each run comprised a learning session (eight pairwise relationships, such as B’→C’ and A→B) followed by a probe session. Note we expanded the original 2×4 sequence to a 2×5 format in order to increase task complexity. During the value learning phase, the final object in each sequence (E or E’) was paired with either a ¥10 or a ¥100 reward icon. Two resting-state sessions (4 minutes each) were included – one before and one after value learning – to mirror the EEG procedure.

Participants again rated their preferences at the end of the task to measure the impact of reward on subjective valuation^42^. A three-way ANOVA, with factors of structural position, time (pre-vs. post-task), and sequence (reward ¥10 vs. reward ¥100) was used to assess impact on final preference ratings. This analysis revealed no significant three-way interaction (*F* (4, 672) = 1.31, *p* = 0.262, *η*² = 0.008; Fig. 6b). However, within each sequence, in either reward magnitude condition, the final object showed a preference increase post-task (reward ¥10: *t* (168) = −2.07, *p* = 0.020; reward ¥100: *t* (168) = −2.34, *p* = 0.010), and preference changes were comparable between these two magnitudes (*t* (168) = −0.18, *p* = 0.858).

When anxiety was considered as a categorically variable (low vs. high, using the same cut-off as in the EEG study), only low-anxiety participants demonstrated a positive linear relationship between proximity to reward and preference changes (reward ¥10: *β* = 0.20, *p* = 0.021; reward ¥100: *β* = 0.18, *p* = 0.029, Fig. 6c). In contrast, high-anxiety individuals did not (reward ¥10: *β* = 0.09, *p* = 0.232; reward ¥100: *β* = −0.12, *p* = 0.130, Fig. 6c). These findings reinforce the conclusion that high trait anxiety disrupts value updating in an offline context, rather than simply reflecting diminished reward sensitivity.

## Discussion

Our study highlights the utility of EEG-based neural decoding for characterizing neural replay, a key process linked to offline learning^13,14^. In line with previous MEG-based replay findings^18,29^, our EEG study revealed a selective increase in backward replay for reward-paired sequences. Moreover, the magnitude of this replay correlated with both behavioural measures of reward learning and a neural index of cognitive map representation. These findings extend previous research in several ways. First, they connect cognitive map representation signatures (offline replay and online RSA) to a specific behavioural disposition – trait anxiety. Specifically, individuals with high trait anxiety displayed reduced reward-related backward replay, which was associated with a distorted neural representation of the reward sequence and a diminished preference shift for reward-predictive stimuli. Further support for this interpretation was evident in a separate behavioural experiment showing inefficient credit assignment among high trait anxiety participants, regardless of reward magnitude. These results align with studies showing that individuals with anxiety have difficulty learning from positive experiences^5,61^ and suggest a potential aberrant replay mechanism during offline learning.

Backward replay has been hypothesised to play a crucial role in offline value learning, by guiding the assignment of value to states and actions within an internal cognitive map^22^. Consequently, we suggest that reduced backward replay in high-trait-anxiety individuals may lead to erroneous value associations or suboptimal credit assignment. Clinically, disrupted connectivity between the hippocampus and the DMN has been reported in anxiety disorders^62–65^, impairing the consolidation of reward-related information and effective decision-making^4,66^. At a more descriptive level, clinical studies have consistently reported reduced hippocampal volumes and heightened hippocampal activation in individuals with anxiety, both clinically and sub-clinically^67–69^.

Rumination, characterised by repetitive, distress-focused thoughts^70,71^, is common in anxiety and depression^72^. Our core finding – that individuals with high trait anxiety exhibit reduced backward replay for rewarded sequences – points to deficits in learning positive outcomes beyond direct associations. Interestingly, during a post-value learning rest period, we observed a marginally significant positive correlation between backward replay for neutral sequences and trait anxiety. This might suggest that individuals with higher anxiety may ruminate on neutral sequences as if they were less favourable experiences. However, this finding should be interpreted with caution, given its modest strength and the neutral (rather than overtly negative) nature of these stimuli. Furthermore, there is little current evidence linking the fast time scale dynamics of replay to elements of conscious experience, as exemplified by rumination. Future research should investigate replay of explicitly negative sequences to elucidate how anxiety shapes the extent and content of neural reactivation, and in relation to rumination^27^.

We did not aim to dissociate trait anxiety from depression, as these overlapping clinical phenotypes may share similar offline learning mechanisms^27^. Indeed, trait anxiety and depression are strongly related: they frequently co-occur in epidemiological studies^49,50^, and trait anxiety is recognised as a biomarker or vulnerability phenotype spanning both conditions^35,36^. In our study, trait anxiety was strongly associated with scores on other mood-related questionnaires, including the SDS, SAS, and PSWQ, implying overlapping clinical characteristics. Rather than focusing on aversive or value-free structure learning, we chose to examine reward learning, given previous work demonstrating the sensitivity of replay to reward^18,19,24^. In future studies, we plan to extend our approach and investigate anticipation of negative outcomes in anxiety^73–75^, as well as the role of aberrant replay in such contexts^3,30^.

Consistent with evidence that trait anxiety reduces learning from positive experiences in online tasks^76,77^, we extend this observation to offline learning by showing that higher trait anxiety coincides with weaker backward replay, diminished neural representation of reward sequences, and smaller preference changes for reward-predictive stimuli. These findings suggest that trait anxiety disrupts replay-based offline value learning beyond potential deficits in structural learning, as indicated by the absence of correlations with structural acquisition or learning efficiency. Future experimental approaches, including longitudinal designs, may help determine whether these relations reflect a single or multiple interacting processes, and refine hypotheses regarding their causal interactions. Additionally, our tasks did not involve structure learning through trial and error, which are processes that other studies^4,78,79^ have implicated in anxiety-related learning deficits. Future research should employ tasks that directly assess structural learning to achieve a more comprehensive understanding of learning deficits in trait anxiety.

Decoding human cognition during rest has considerable promise for elucidating mechanisms underpinning learning, memory, and planning during offline periods^12^, and thereby offer fresh perspectives on psychiatric disorders^1,80,81^. Such work has driven methodological developments in neural decoding and replay analysis using non-invasive neuroimaging^18,28,33^. MEG studies, in particular, offer high spatiotemporal resolution and have demonstrated replay during rest^1,3,29,30,32^. By comparison, EEG offers comparable temporal resolution and has the added benefit of being more readily available in clinical settings. The current study demonstrates that a 64-channel EEG system can decode up to eight stimuli with sufficient sensitivity to detect their sequential neural replay during rest periods (Supplementary Fig. 6). Consistent with prior MEG research, we observed a selective increase in backward replay for reward sequences following value learning, but no replay for visual sequences.

Unlike previous MEG studies^1,18^, we did not detect forward replay prior to value learning. This may be due to our subject selection procedure, which excluded participants who failed the Day-1 structural learning requirement, thereby ensuring that structural knowledge differences did not confound value learning in individuals with varying trait anxiety. However, this approach contrasts with that of Nour et al.^1^, who implemented extensive training to avoid excluding participants. Consequently, in the current study, skilled in structural learning may have less reliance on replay during rest, potentially leading to a reduced need for offline structure learning ^39–41^. Future research can conider alternative designs that enable a more sensitive assessment of how replay is influenced by task performance. Furthermore, while 64-channel EEG systems were adequate for detecting replay (see also Supplementary Fig. 6), the lower decoding accuracy (mean accuracy 23.21 ± 0.90%) compared to MEG (275 independent sensors and mean accuracy 39.60 ± 2.20%)^18^, might further limit replay detection. Future work may benefit from improved EEG decoding approaches, e.g., spectral frequency decomposition^82,83^ and high-density configurations^84,85^, both of which might enhance replay detection.

Another difference in the present study is that it was conducted with a Chinese-speaking sample. Nonetheless, the use of non-linguistic images minimises potential language-based influences on replay, and we have successfully detected neural replay in a similar population with simultaneous EEG–fMRI recordings^86^. Preliminary analyses indicate that these findings align with those from other populations, suggesting that the fundamental neural mechanisms for offline learning are broadly conserved across cultural and linguistic backgrounds^1,18,29^.

In conclusion, our findings demonstrate that replay measurement is feasible using clinically accessible EEG technology, highlighting the importance of considering offline learning mechanisms in neuropsychiatric disorders. We show that high trait anxiety correlates with aberrant offline learning signatures, specifically reduced backward replay for reward sequences during rest – an impairment associated with altered cognitive map representation both neurally and behaviourally. The approach we describe opens new avenues for investigating offline learning mechanisms in psychopathology, including potential interventions that can modulate memory and affective processes during rest or sleep.

## Methods

### Participants

#### EEG experiment

Eighty healthy volunteers (46 males; mean age = 20.90 ± 1.95) participated in the EEG experiment. To achieve a statistical power of 0.80 for detecting the effect size (0.35) based on previous similar studies^4,87,88^, around 60 participants should be recruited. Considering some participants may not be able to pass the Day-1 structure learning, we expanded our sample size to 80 individuals. All participants had a normal or corrected-to-normal vision and no history of neurological or psychiatric illness. The study was approved by the local Ethical Committee of Shenzhen University (PN-202300013).

Participants were recruited based on their trait anxiety score on the Chinese-validated version of STAI^37,38,59,60^ so that they evenly spanned its full spectrum (20 subjects in four score domains, 20-30, 30-40, 40-50, and above 50). The overall mean trait anxiety score was 42.57 ± 1.34. The participants provided written informed consent in compliance with the Declaration of Helsinki and were compensated approximately ¥200 (¥100 show-up fee plus a variable amount up to ¥100 depending on task performance) for their participation. The participants received comprehensive training on the task rule (structure learning) one day prior to the EEG task and only subjects achieving at least 80% accuracy were permitted to the EEG experiment the next day. Seven participants were excluded from the analysis because of failure of passing the Day-1 structure learning, and another five subjects were excluded due to excessive movements and/or recording artifacts during EEG recording. A total of 68 subjects were included for subsequent analysis.

The trait anxiety was treated as a continuous variable in all relevant analyses. In a complementary analysis, we also found consistent findings in group analysis (see Supplementary materials) based on the normative trait anxiety scores for the Chinese population^38^: those who scored 45 or higher were assigned to the high trait anxiety group (32 subjects, mean score = 53.41 ± 6.63), while those who scored below 45 were placed in the low trait anxiety group (36 subjects, mean score = 35.50 ± 6.01). Previous studies indicate that individuals with high trait anxiety (≥ 45) exhibit behavioural and neural deficits similar to those with anxiety disorders^51,89–91^.

#### Behavioural experiment manipulating reward magnitude

Two hundred healthy volunteers participated in a separate behavioural experiment, all of whom had normal or corrected-to-normal vision and no history of neurological or psychiatric illness. Thirty-one of these were excluded from the analysis for failing to pass the structure learning task or completing the whole task, resulting in a final sample of 169 subjects (106 females; mean age = 19.89 ± 2.03). Participants were again recruited based on their trait anxiety scores from the Chinese-validated version of STAI^37,38,59,60^, yielding an overall mean trait anxiety score of 44.68 ± 8.78. Those scoring 45 or higher were assigned to the high trait anxiety group (87 subjects, mean score = 51.68 ± 5.21), while those scoring below 45 were assigned to the low trait anxiety group (82 subjects, mean score = 37.24 ± 4.77)^37,38^. All participants provided written informed consent in accordance with the Declaration of Helsinki and were compensated approximately ¥60 (¥30 show-up fee plus up to ¥30 based on task performance).

#### Design and Materials

##### EEG experiment

The task spanned two days in a manner that followed closely our previous MEG study design^18^, with the exception of eliciting stimuli preference ratings before and after the main task, in order to capture a putative reward effect on behaviour.

On Day-1, participants were shown eight sample stimuli embedded in two visual sequences [B’/A/D’/B] and [A’/C/C’/D] and were explicitly instructed on how to map the stimuli onto two true sequences: A->B->C->D and A’->B’->C’->D’. This mapping between visual and underlying true order was randomised across subjects. Subjects completed four runs of training where each run consisted of one learning and one probe session. During learning, two visual sequences ([B’/A/D’/B] and [A’/C/C’/D]) were presented three times, with each stimulus presented serially in the centre of the screen for 900 ms, followed by a 500 ms inter-stimulus interval. Each sequence presentation ends with a 2000 ms interval of blank screen. After that, participants were quizzed about the true order of the stimuli without feedback. On each trial, the target image appeared for 5000 ms, during which subjects were required to think about which image should appear following (but not necessarily next to) the target image. When considering directional (i.e., forward) sequences, we distinguish between a picture that immediately follows the target (“next to” or “adjacent,” e.g., 1→2) and a picture that appears at any point after the target (“following,” e.g., 1→4). The former is a subset of the latter. This distinction is crucial for assessing whether participants have accurately learned, and can subsequently reproduce, the correct order of the true sequence^1,18^. Then, two candidate images appear on the screen and subjects were asked to select the correct one within 1000 ms, a limit set to minimise potential associative learning when the stimuli were presented together^18^. Participants were required to reach at least 80% accuracy in the last structure learning run to be permitted for the Day-2 experiment.

On Day 2, participants were presented with a new set of eight stimuli, and first rated their preferences for each of the eight images on a 1-9 scale (from “strongly dislike” to “strongly like”). The procedure was performed twice, one before and one after the main task, where the aim was to assess preference changes due to value learning^42,92^, an implicit reward effect on behaviour. Subjects then performed the main task during concurrent whole-brain EEG recording. The main task comprises four phases: functional localiser, sequence learning, value learning, and a position test, and three resting states, one immediately before the functional localiser task, one before and one after value learning in order to capture reward induced change in spontaneous neural replay.

In the functional localiser phase, stimuli were presented in a random order. As this preceded sequence learning, participants had no structure information about stimuli, such that decoding models trained during functional localiser solely captured sensory level stimulus information. On each trial, this phase starts with a text presented for 1500 - 3000 ms during which subjects were required to vividly imagine its associated imagery content, followed then with the actual image for 750 ms, and a jitter ITI for 700 - 1700 ms. To ensure subjects were attending to the stimuli, on rare occasions (20% probability), random stimuli were presented upside down and subjects were required to respond to these by pressing a button. There were 400 trials in total, split in two runs, with 50 instances for each visual stimulus (20% were upside down). Only correct-side-up images (40 repetitions per image) were used for classifier training.

In the sequence learning phase, subjects were required to apply the Day-1 learnt mapping rule to the Day-2 new stimuli. The use of an entirely new set of stimuli ensured neural replay signatures in EEG (if exist) could not be attributed to perceptual biases introduced during Day-1 training. Participants were explicitly informed that the mapping rule was the same with the one on Day-1. There were three runs of sequence learning. As for structure learning, each run consisted of one learning and one probe session. During learning, two visual sequences ([B’/A/D’/B] and [A’/C/C’/D]) were presented three times, with each stimulus presented serially in the centre of the screen for 900 ms, followed by a 500 ms inter-stimulus interval. Each sequence presentation ends with a 2000 ms interval of blank screen. After that, participants were quizzed about the true order of the stimuli without feedback. On each trial, the target image appeared for 5000 ms, during which subjects were required to think about which image should appear following (but not necessarily next to) the target image. Then, two candidate images were presented on the screen and subjects were tasked to select the correct one within 1000 ms. This time limit was designed to minimise potential associative learning when the stimuli were presented together. No feedback was provided. There was a 33% possibility that the wrong answer came from the same sequence but preceded instead of following the probe stimuli. This setup was designed to encourage participants to form sequential rather than clustering representations (i.e., which sequence does this object belong to).

Next, participants completed value learning, in which they were taught that the end stimulus of one of the sequences would lead to monetary reward, while the end stimulus of the other would not, in a deterministic way. Specifically, on each trial, participants saw the object at each end of the sequence (i.e., D or D’) for 900 ms, followed by an inter-stimulus interval of 3000 ms, and then either received a reward (image of 10 Chinese Yuan banknote) or no-reward (scrambled image of 10 Chinese Yuan banknote) outcome for 2000 ms, followed by an ITI of 3000 ms. Participants were required to press one button for the reward and a different button for non-reward. Pressing the correct button to “pick up” the coin led to a payout of this amount at the end of the experiment, and participants were pre-informed of this. There were 12 repetitions for each association, with trial order randomised. Before or after the value learning, participants underwent a 4-minute resting session.

Participants subsequently completed a position test, wherein an object was presented on screen for 1000 ms, and subjects required to think about its position within its sequence. Each object was followed by a single number (1, 2, 3, or 4), and participants indicated whether the presented number matched the sequence position of the preceding object, via a button press (1000 ms response window, chance accuracy 50%). This was followed by a 3000 ms ITI. There were 80 trials in total, 10 repetitions for each stimulus, with a constraint that the same stimulus does not appear consecutively. No feedback was provided. Finally, at the end of the main task, participants were required to write down the true sequences in the correct order.

Notably, participants did not receive any direct monetary rewards on Day 2 for identifying correct positions, as the focus was on offline value learning and replay rather than immediate rewards. The ≥ 80% accuracy criterion on Day 1 nonetheless ensured that all participants had already learned the abstracted task structure.

##### Behavioural experiment manipulating reward magnitude

This behavioural study comprised six phases: pre-preference rating, structure learning, a rest period before value learning, value learning, a rest period after value learning, and post-preference rating. To evaluate how value learning impacted subjective valuation, participants began by rating their preferences for ten stimuli on a 1–9 scale, both before the main task and after completing it.

During the structure learning phase, participants were required to learn pairwise rank relationships between the objects to infer the true sequences, specifically [A/B/C/D/E] and [A’/B’/C’/D’/E’]. Three runs of sequence learning were conducted, each containing one learning and one probe session. During learning, eight pairwise image combinations were displayed three times each (1500 ms per stimulus, 500 ms ISI). This was followed by a quiz session (without feedback) encouraging sequential rather than cluster-based learning.

In the value learning phase, the final object of each sequence (E or E’) was paired with either a ¥10 or ¥100 reward icon (a real banknote image). Each trial displayed the object for 900 ms, followed by a 1000 ms ISI, then the reward image for 2000 ms, followed by a 1000 ms ITI. Participants pressed different buttons to register whether the stimulus was associated with a ¥10 or ¥100 reward and could earn that amount (divided by a constant factor) by responding correctly. Each association was presented 12 times in random order. Resting periods (4 minutes each) occurred before and after value learning.

Finally, participants performed a post-task preference rating to assess changes in preferences following value learning. Although offline replay could not be directly measured in this behaviour experiment, these rest sessions were designed to parallel the EEG study, implying that any offline replay might also occur during these intervals. This design enabled us to examine whether reward magnitude influenced behavioural responses, offering insights into the underlying mechanisms of offline value learning in sequences.

## EEG Data Acquisition

### Data Acquisition and Preprocessing

In this study, a Brain Products 64 channel electroencephalography (EEG) system was used to record EEG during Day-2 experiment, with an additional electrode placed below the right eye as vertical electro-oculograms (EOG). Online EEG recordings were sampled at a rate of 1000 Hz with FCz as reference electrode during data acquisition. All electrode impedances were kept below 5 *K*Ω.

Preprocessing was conducted separately for each session, following the approach used by previous MEG study^18^. The data were first high-pass filtered at 0.5 Hz and notch filter of 50 Hz to remove line noise, and then down-sampled to 100 Hz. The continuous recording was visually screened for noisy channels, which were interpolated using the weighted average of surrounding electrodes. All electrodes were re-referenced offline to the average. The data was subsequently segmented into epochs, with each epoch spanning from-100 ms before to 750 ms after the image onset. Trials with extremely high noises by visual inspection were manually excluded. Bad channels with abnormal voltage were interpolated with the weighted average of their neighboring electrodes. Independent component analysis (ICA) was performed to remove eye movement and other artifactual components and the remaining components were then back projected to the EEG channel space. Five participants were excluded for excessive head movement or other artifacts. Only trials with correct button press were used for subsequent analyses.

## EEG Analysis

### Neural Decoding Analysis

Same with Liu et al.^18^, we trained each of the eight binary classifiers (*k* ∈ {1:8}) on the evoked whole-brain neural response at a signal time bin after the stimulus onset in the functional localiser task. Essentially, model (*k*) distinguished sensor patterns based on the particular stimulus *k* (positive examples) relative to all other stimuli (∼*k*), which was further aided by an equal amount of “null” data from the inter-trial interval that acted as negative examples. The inclusion of null data decreased the spatial correlation between the decoders and allowed simultaneous reporting of low probabilities in the rest data by all decoders^18^. To enhance sensitivity for sequence detection, we used L1 regularization and fixed lambda = 0.001, to prevent overfitting of the results to the regularization parameter. Every model *k*, unique to time and stimulus, was represented as a single vector of length *n* + 1 (*n* sensors [max. 62], plus intercept).

To quantify within-session decoding accuracy, we trained regression models using 10-fold cross-validation on the functional localiser data. During each validation loop, prediction accuracy was quantified as the proportion of test trials, where the decoder reported the highest probability corresponding to the trial label, giving a prediction accuracy. Each classifier is trained to recognise only one of the eight images, the probability of correctly identifying an image in a balanced dataset defaults to 1/8 (i.e., 0.125), which represents chance performance. The average of all estimates obtained over the validation loops determined the overall accuracy, consistent with the approach in previous MEG studies^1,92^.

Decoders trained and tested at 180 ms post-stimulus onset achieved the highest decoding accuracy (mean accuracy across participants; Fig. 2a and Fig. S1). Accordingly, these decoders were employed in the replay analysis^18,31,93^. We confirmed that decoding accuracy was significantly above chance (12.5%) using nonparametric tests. The association between the eight visual stimuli and their respective state indices was fixed within each participant but randomised across participants, ensuring that any stimulus-related factor (e.g., overall preference or decodability) did not systematically affect state decoding at the group level. During the functional localiser phase, participants received no information about stimulus–state mappings and the images were presented in random order, so classifiers remained free of task-related biases. We also trained all classifiers exclusively on pre-learning EEG data to avoid contamination by task-specific sequence knowledge^28^, and then applied them to the resting-state EEG to examine replay patterns.

### Neural Sequence Analysis

To evaluate neural sequences, we measured the sequence strength of a pairwise state-to-state transition using a multiple regression model, where the representation of state *i* statistically predicts the subsequent representation of state *j* at a specific time lag (i.e., speed of replay). This measure is an average estimate of statistical predictiveness, incorporating the number and strength of replay events, and is referred to as “sequence strength.” The reason for this approach is that neural representations (of different states) are efficiently decoded in a noisy and probabilistic manner. The detailed methodology, including related simulations, is documented in Liu et al.^18^. We have implemented the Temporal Delayed Linear Modelling (TDLM) methodology in EEG data following its success in earlier MEG empirical work. The process began by utilizing eight-state classifiers (from the highest accuracy time bin) to characterise EEG data from each timepoint of all the resting sessions, thus generating a [time x state] reactivation probability matrix for each session. We then utilised TDLM to estimate the evidence for sequential reactivations, consistent with the task-defined sequential order.

TDLM is a method that employs multiple linear regression to quantify the degree to which a lagged reactivation time course of state *i*, (*X*(Δ*t*).), where *t* indicates lag time) can predict the reactivation time-course of state *j*, (*X*_0_). This approach comprises two stages. Initially, we conducted separate family multiple regressions (first-stage) with the reactivation time course of each state (*j* ∈ {1:8}) as the dependent variable and the historical (i.e., time-lagged) reactivation time courses of all states as predictor variables:

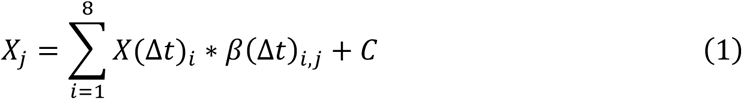

The predictor (design) matrix from a single model contained a separate predictor for the reactivation time courses of all states (*i* ∈ {1: 8}), lagged by Δ*t* ∈ {10 ms, 20 ms,…, 90 ms}, plus the reactivation time course of all states lagged by Δ*t* + α, where α = [100 ms, 200 ms,…], to capture autocorrelations in state time courses at a canonical alpha frequency, which predominated in human brain activity at rest, in addition to a constant term, *C*. We repeated the regression in Equation 1 for each *j* ∈ {1: 8} and Δ*t* ∈ {10, 20, 30,…, 90 ms}, and used ordinary least squares regression to obtain β. The regression coefficients from Equation 1 quantify the evidence for each empirical state-to-state reactivation pattern at a specific lag. Δ*t*. For example, β(Δ*t*)_.,0_ quantifies the coefficient that captures the unique variance in *X*_0_ explained by *X*(Δ*t*). These coefficients were demonstrated in a lag-specific [8 x 8] empirical transition matrix ***B***, representing evidence for all possible state-to-state transitions at a given time lag.

In a second-level regression, we quantified the evidence that the empirical transition matrix, ***B***, can be predicted by the underlying task transition structure (i.e., true sequences).

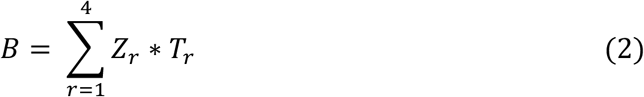

Here, ***T***_r_ is the [state x state] predictor transition matrix (for regressor *r*, with 1 for transitions of interest, and 0 otherwise), and ***Z***_r_ is the scalar regression coefficient quantifying the evidence that the hypothesised transitions, ***T***_r_ predict the empirical transitions, ***B***. Four predictor matrices were considered, including ***T**_auto_*, ***T**_const_*, ***T**_F_*, ***T**_B_*, (1) ***T**_F_*: true sequence transitions in the *forward* direction (A->B->C->D and A’->B’->C->D’ entries in ***T**_F_* = 1, all others set to 0), (2) ***T**_B_*, true sequence transitions in the *backward* direction (D->C->B->A and D’->C’->B’->A’, i.e. ***T**_B_* is the transpose of ***T**_F_*), (3) ***T**_auto_*: self-transitions ([8 x 8] identity matrix), to control for auto-correlation, and (4) ***T**_const_*: a constant matrix, to capture the average of all transitions, ensuring that any weight on ***T**_F_* and ***T**_B_* was not due to general dynamics in background neural dynamics.

***Z*** represents the weights of the second-level regression, which is a vector with dimension of *r* by 1. Each entry in ***Z*** reflects evidence for the hypothesised sequences in the empirical transitions, i.e., sequence strength. Note that this estimate of sequence strength is a relative quantity, and an estimate of zero for state *i* to state *j* does not mean there is no replay of *i*-> *j*, but rather suggests that there is no stronger replay of *i*-> *j* than that of other transitions.

The regression in Equation 2 was repeated for each time lag (Δ*t* = 10, 20, 30,…, 600 ms), resulting in time courses of both forward and backward sequence strength as a function of time lag. Shorter lags indicate greater time compression, corresponding to faster speed. For statistical testing, we used non-parametric permutation tests at the second-level regression, shuffling the rows and columns of ***T***_’_ (*forward* predictor matrix), defining ***T**_B_*: (*backward* predictor matrix) as its transpose. For each of 100 permutations we calculated the peak absolute mean sequence strength over participants and across lags (controlling for multiple comparisons across lags). Sequence strength in the unpermuted data was deemed significant at peak-level *P*_FWE_ < 0.05 if its absolute magnitude exceeded 95% of within-permutation peak.

### Representational Similarity Analysis (RSA)

To examine how the representation of task structure change from the functional localiser (pre-learning) to Position Probe (post-learning) sessions, we applied RSA on EEG data. We first z-scored the pre-processed EEG data across all trials for each sensor and time point (*t*,-100 to +750 ms) post stimulus-onset. Next, we regressed the [trial x 1] neural data for each trial, *Y(s)_t_* (at time point *t* and sensor *s*), onto a session design matrix, **X**, which included dummy coding for the trial stimulus label, using Equation 3:

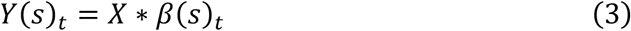

Here, β(*s*)*_t_* was the [stimulus x 1] vector of regression weights, as an estimate of the stimulus-specific activation for sensor *s* at time point *t*. The procedure was repeated over all sensors to yield a [sensor x stimulus] matrix at each time point. Pearson correlation distance was computed between the sensor patterns for each pair of pictures (columns), which resulted in a symmetrical [8 x 8] Representational Dissimilarity Matrix (RDM) at each time point. This procedure was repeated in functional localiser (FL) and Position Probe (PP) sessions to determine the learning-induced increase in representational similarity Δ***S**_t_* at each time point, as expressed in Equation 4:

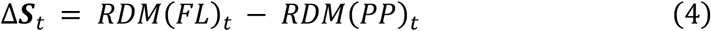

Here, *s*_.0_ of Δ***S**_t_* quantified the post-learning similarity increase between evoked signals for stimuli *i* and *j* at time *t*.

To assess the unique contribution of three predictors (an abstracted representation of ordinal position, reward within-sequence, and neutral within-sequence representation) in explaining the variance in Δ***S**_t_*, we employed a second multiple regressions approach. The data were preprocessed by smoothing Δ***S**_t_* over time using a 90 ms Gaussian kernel before conducting regression analysis. To identify time windows (clusters) that showed significant positive evidence for each predictor, we utilised nonparametric tests while correcting for multiple comparisons over time. Specifically, we performed correlation analysis with subjects’ trait anxiety for each time point over all participants for each predictor and computed the sum of *r*-values within each continuous stretch of time points exhibiting a positive effect at α level of 0.05. And we also performed one-sample *t*-tests for each time point over all participants for each predictor and computed the sum of *t*-values within each continuous stretch of time points exhibiting a positive effect at α level of 0.05.

To validate our results, we repeated this procedure 1000 times, shuffling the rows and columns (stimulus labels) of Δ***S**_t_* consistently across time to preserve temporal smoothness prior to the second regression (permutations). Finally, we extracted the maximal sum-of-*r* or *t* value for each predictor and identified a suprathreshold cluster in the unpermuted data as significant if its sum-of-*t* exceeded 95% of maximal within-permutation sum-of-*t* values. This method is the same with previous study^1^. The results are displayed in Fig. 5b.

## Acknowledgment

Conceptualization, Y.L., Q.Y., R.D. and Y.Luo.; Investigation, Q.Y., Y.L., C.H., J.O., H.W., Z.X.; Writing – Original Draft, Q.Y., Y.L.; Writing – Review & Editing, Q.Y., Y.L., M.N, and R.D. This study is supported by the National Science and Technology Innovation 2030 Major Program (2022ZD0205500), the National Natural Science Foundation of China (32271093, 31920103009), the Beijing Natural Science Foundation (Z230010, L222033), the Fundamental Research Funds for the Central Universities, the Major Project of National Social Science Foundation (20&ZD153) and Shenzhen-Hong Kong Institute of Brain Science – Shenzhen Fundamental Research Institutions (2022SHIBS0003).

## Data & code availability

Data and code will be openly shared on publication. All analysis scripts will be released at https://gitlab.com/liu_lab/EEGReplay.git. The data will be deposited on the Open Science Framework (OSF) at https://osf.io/gjvsc/ and https://osf.io/j34hw/. Source data for every figure will accompany the paper.

## Supplementary Information

**Table 1:** The Chinese-validated version of Spielberger Trait Anxiety Inventory (STAI) Questionnaire (the Chinese version and English version).

| 题目 | 完全没有 | 有些 | 经常 | 几乎总是如此 |
| --- | --- | --- | --- | --- |
| 我感到愉快 | 1 | 2 | 3 | 4 |
| 我感到神经过敏和不安 | 1 | 2 | 3 | 4 |
| 我感到自我满足 | 1 | 2 | 3 | 4 |
| 我希望能像别人那样高兴 | 1 | 2 | 3 | 4 |
| 我感到我像衰竭一样 | 1 | 2 | 3 | 4 |
| 我感到很宁静 | 1 | 2 | 3 | 4 |
| 我是平静的、冷静的和泰然自若的 | 1 | 2 | 3 | 4 |
| 我感到困难一一堆集起来，因此无法克服 | 1 | 2 | 3 | 4 |
| 我过分忧虑一些事，实际这些事无关紧要 | 1 | 2 | 3 | 4 |
| 我是高兴的 | 1 | 2 | 3 | 4 |
| 我的思想处于混乱状态 | 1 | 2 | 3 | 4 |
| 我缺乏自信心 | 1 | 2 | 3 | 4 |
| 我感到安全 | 1 | 2 | 3 | 4 |
| 我容易做出决断 | 1 | 2 | 3 | 4 |
| 我感到不合适 | 1 | 2 | 3 | 4 |
| 我是满足的 | 1 | 2 | 3 | 4 |
| 一些不重要的思想总缠绕着我，并打扰我 | 1 | 2 | 3 | 4 |
| 我产生的沮丧是如此强烈，以致我不能从思想中排除它们 | 1 | 2 | 3 | 4 |
| 我是一个镇定的人 | 1 | 2 | 3 | 4 |
| 当我考虑我目前的事情和利益时，我就陷入紧张状态 | 1 | 2 | 3 | 4 |

### The Spielberger Trait Anxiety Inventory (STAI, English version) Instructions

Listed below are some of the statements that people often use to describe themselves. Read each statement and put a “√” under the appropriate number on the right to indicate your most appropriate feeling in general, that is, the most appropriate feeling you have often. There is no right or wrong answer. Don’t spend too much time thinking about any one statement.

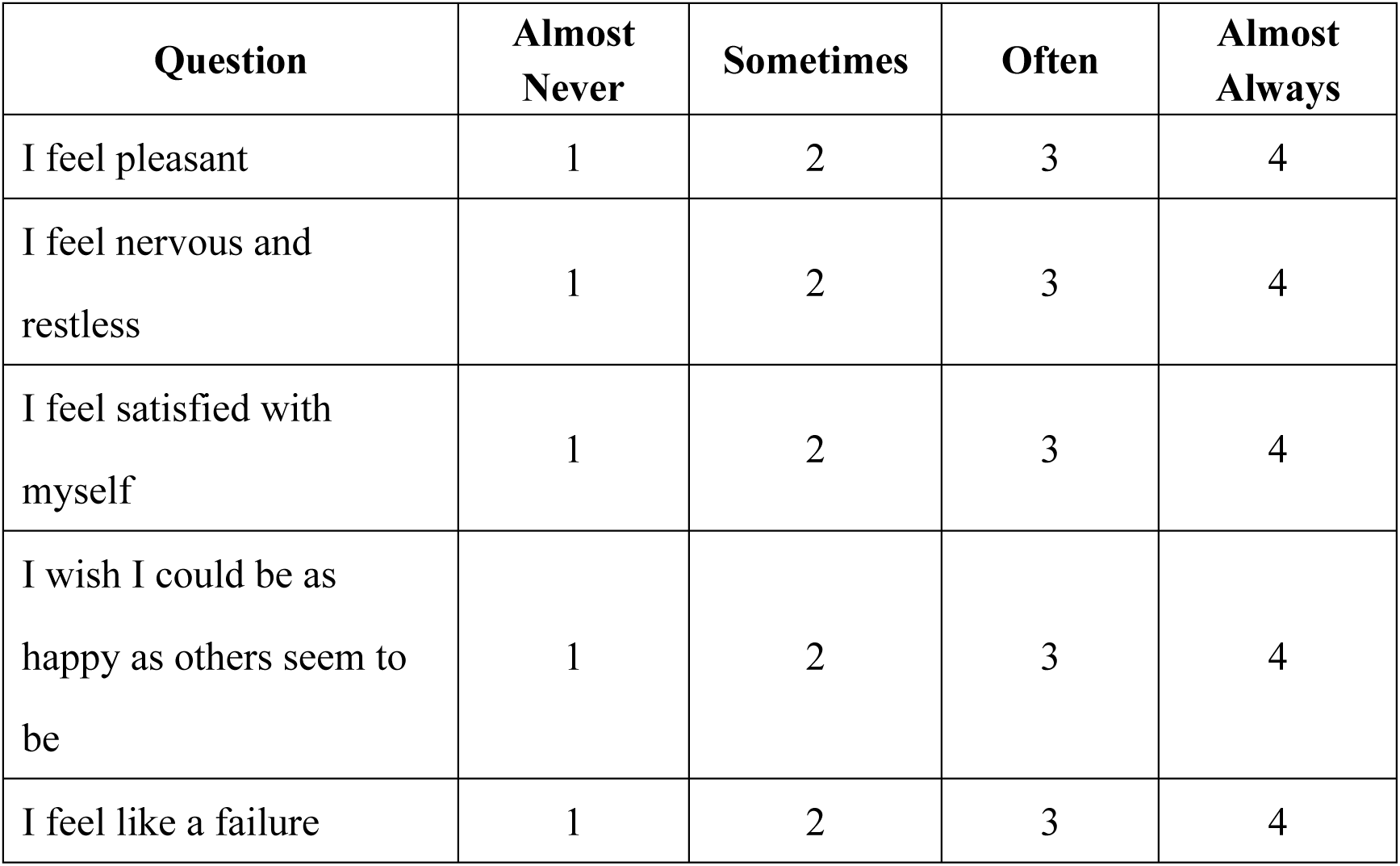
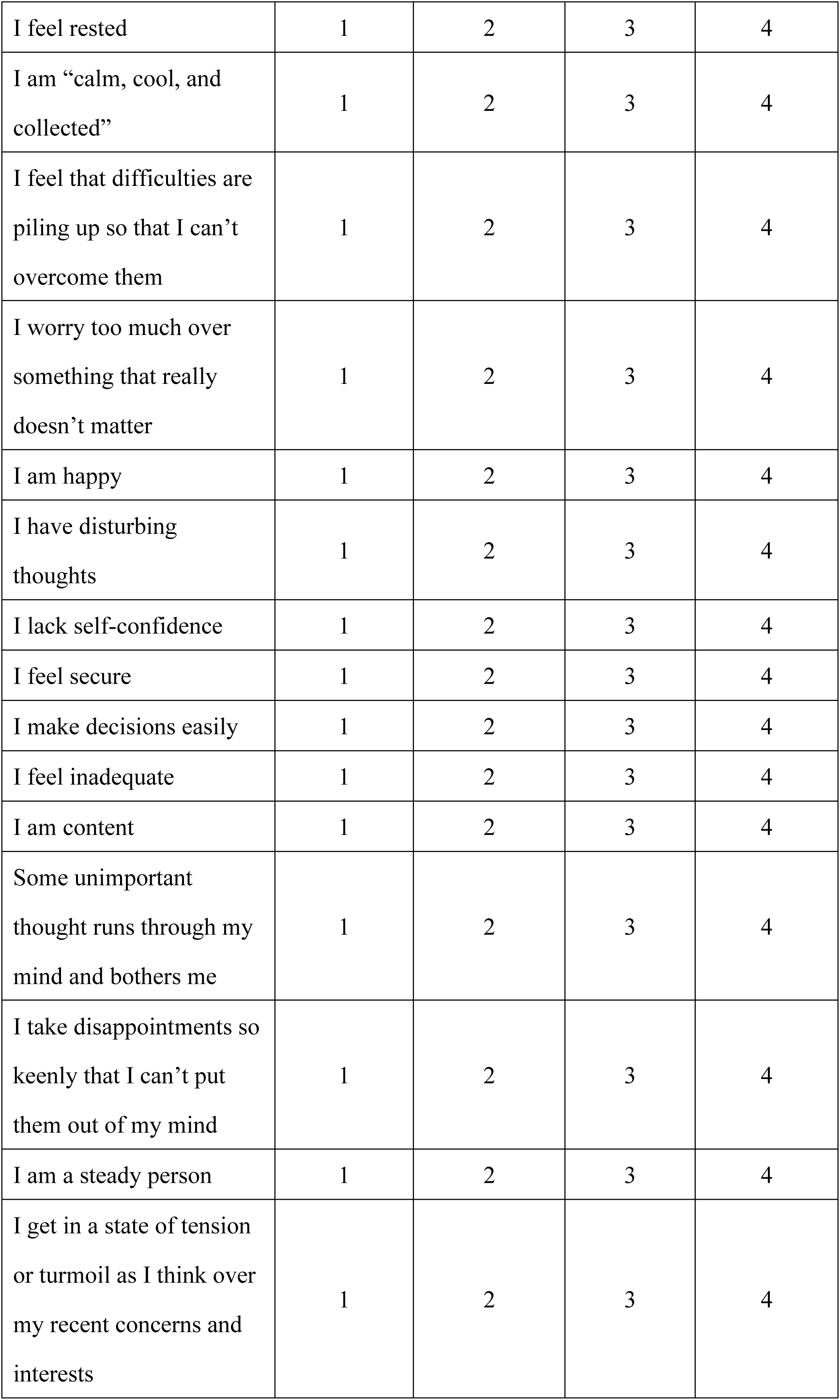

---

**Supplementary Fig. 1.**
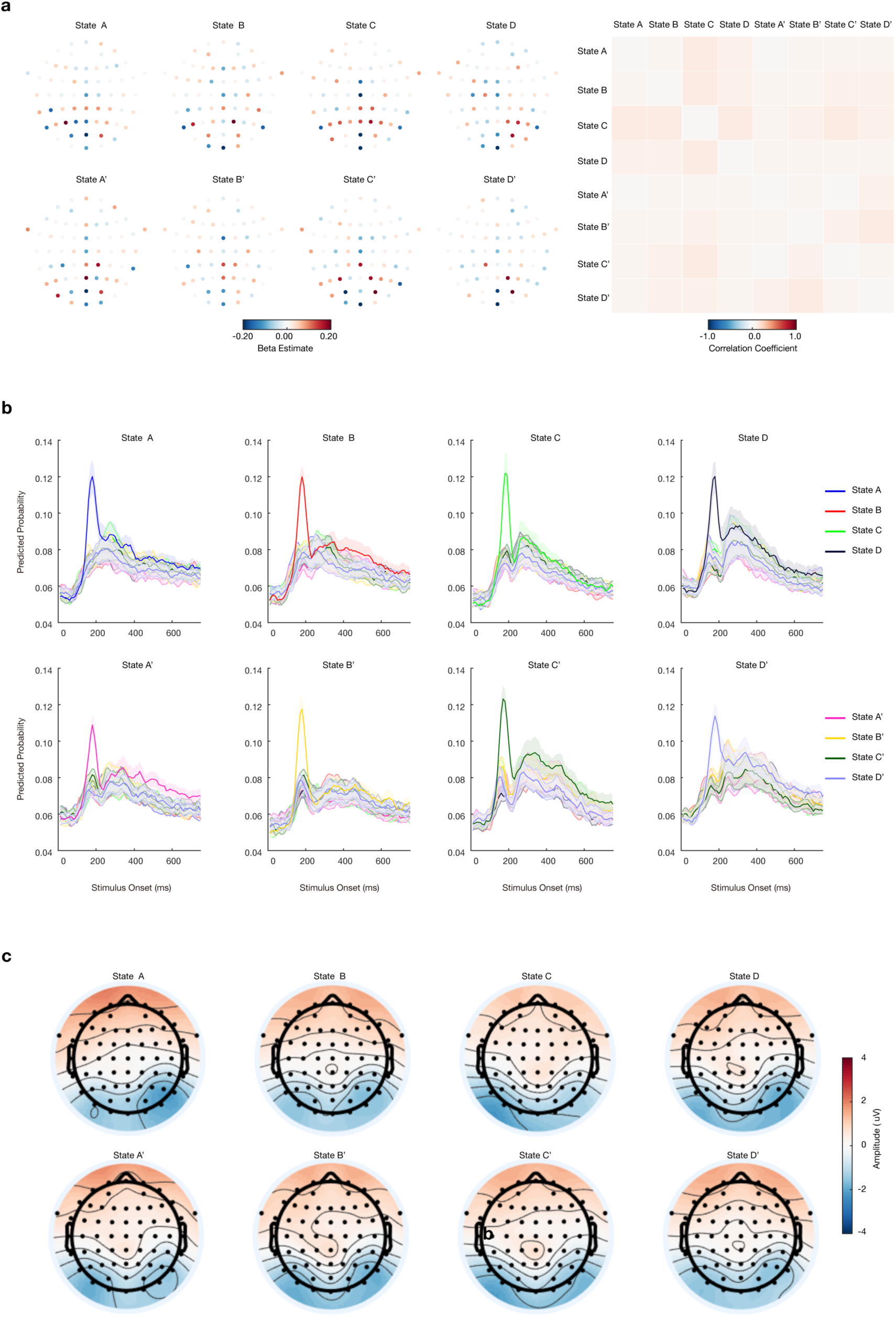
Sensor Maps, Spatial Correlation, and Classifiers Performance of Trained Lasso Logistic Regression Models, Scalp map, Related to. Figure 2**. a**, Sensor map for each state decoding model is shown on the left, with correlation matrix between classifiers shown on the right. **b**, leave-one-out cross-validation results for each classifier in functional localiser task. These plots only use classifiers trained at 180 ms post stimulus onset. The x axis refers to the time point used for testing the classifiers. The target state of the decoder is indicated by line color, and the ground truth state is represented by a different panel for each state. The curves therefore have a different shape than plots made by varying both the training and testing time. **c**, Scalp map show response amplitude for each state at 180ms post stimulus onset.

**Supplementary Fig. 2.**
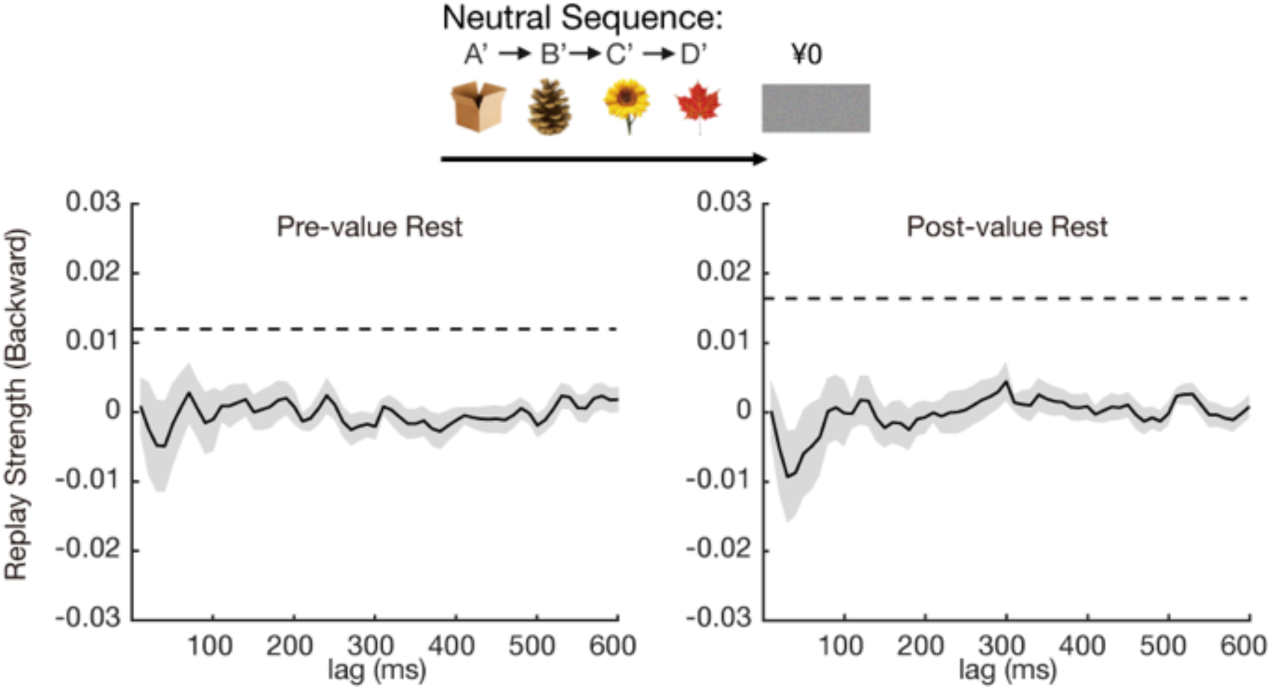
Backward Replay of Neutral Sequences, Before and After Value Learning, related to. Figure 3. No evidence of backward replay of neutral sequence. The dotted line is the FWE-corrected significance threshold.

**Supplementary Fig. 3.**
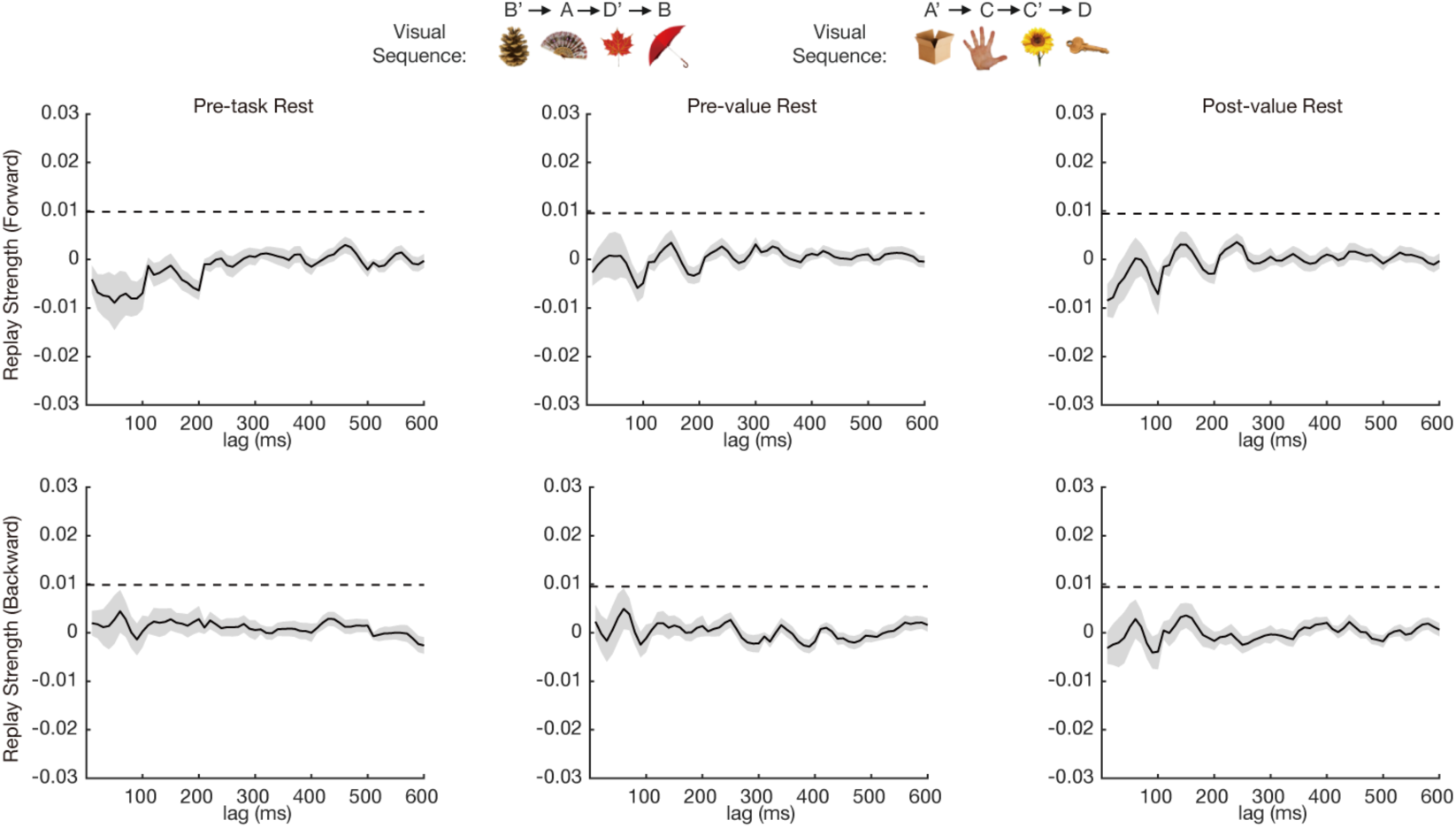
No Evidence of Replay of Visual Sequences, related to. Figure 3. There was no evidence for replay of the visually experienced sequence during 3 resting states among the whole task, same with previous MEG studies. The dotted line is the FWE-corrected significance threshold.

**Supplementary Fig. 4.**
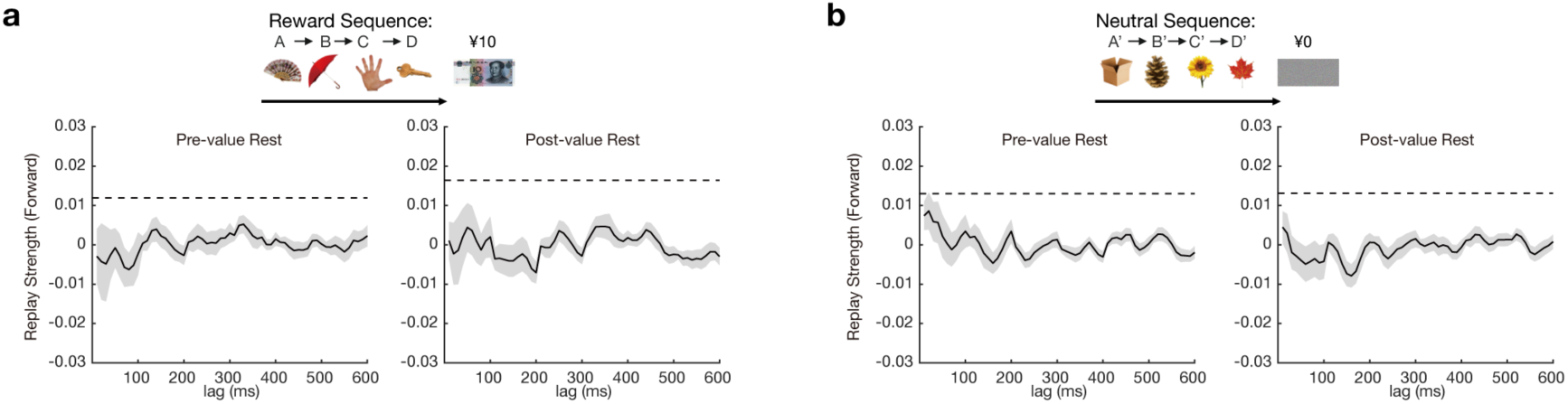
Forward Replay of True Sequences, Before and After Value Learning, related to. Figure 3. No evidence of forward replay of sequences. The dotted line is the FWE-corrected significance threshold.

**Supplementary Fig. 5.**
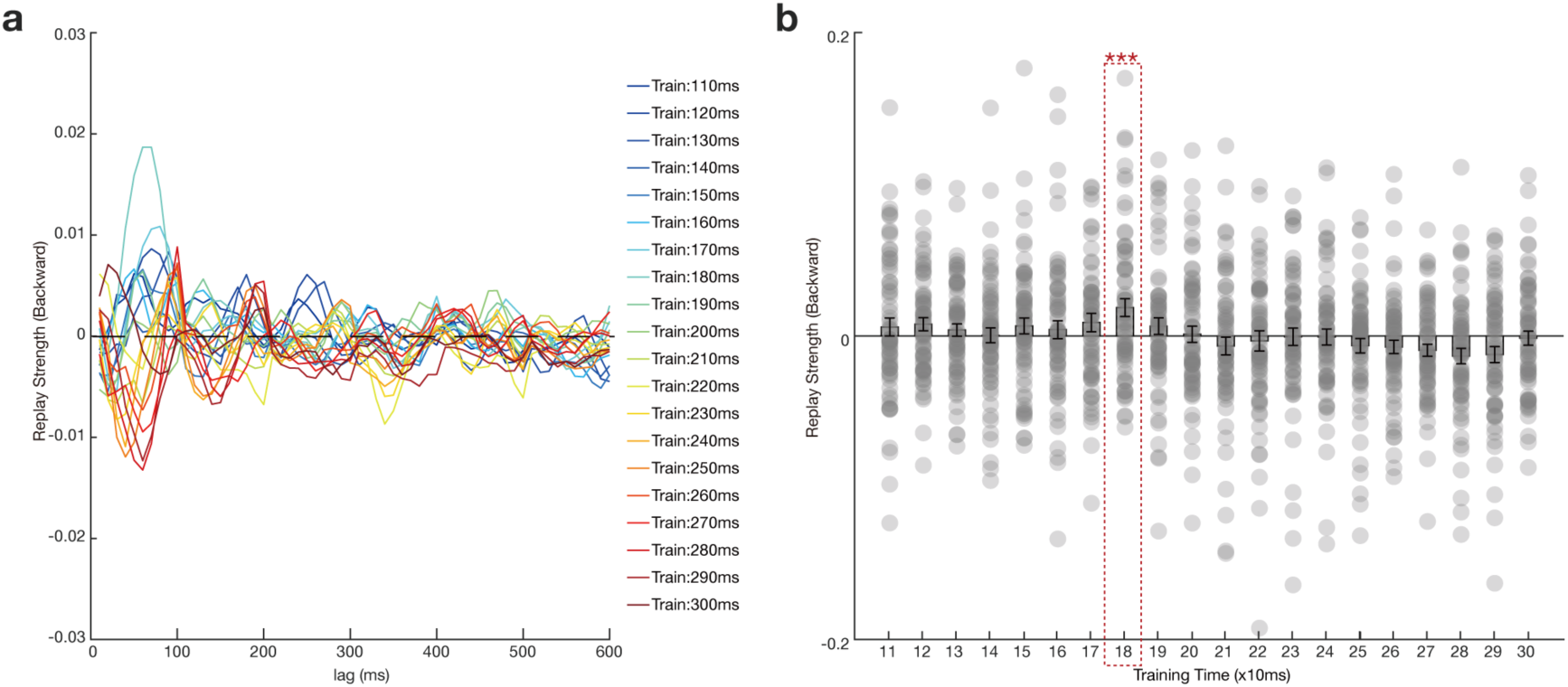
Replay Strength Analysis with Classifiers Trained at Different Times, Related to. Figures 3**. a,** Replay strength using classifiers trained at different times relative to stimulus onset (110 ms – 300 ms). This is from resting data after value learning. **b,** Scatterplot of the replay strength at 60 ms time lag as a function of classifier training times during rest period after value learning. 180 ms is the training time used throughout the current study. *** is *p* < 0.001.

**Supplementary Fig. 6.**
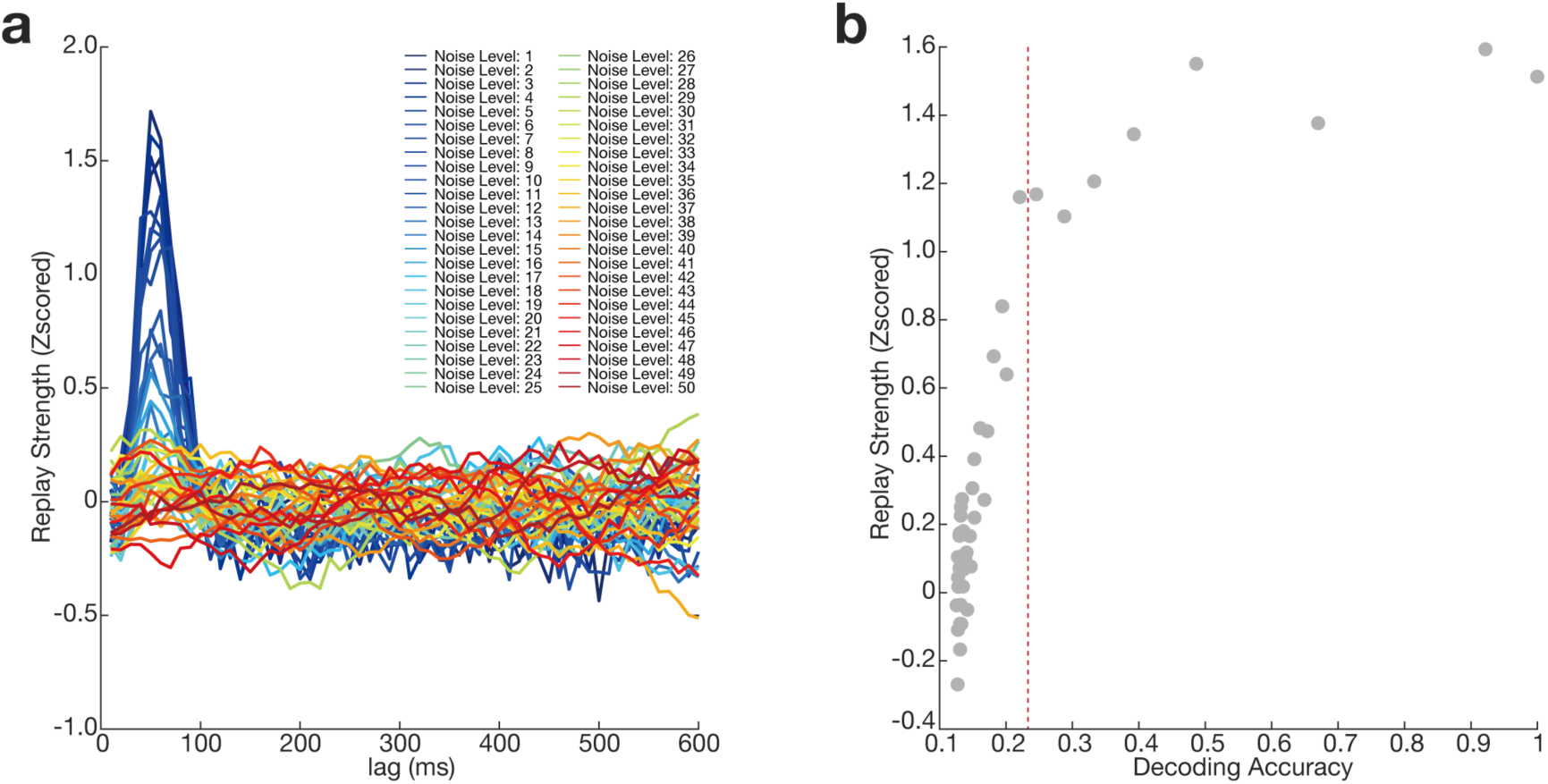
Simulation Results for Decoding Accuracy and Replay Strength, Related to. Figures 3. In the simulation we varied decoding accuracy by injecting different levels of noise during classifier training. We then ask how does decoding accuracy influence replay strength (fixed and known in simulation) using current sequence detection methods. **a,** Sequence strengths were plotted for a classifier trained with different noise levels. **b,** The sequence strength at 60 ms lag (ground truth) is plotted against corresponding decoding accuracy. The red dotted line indicates the empirical decoding accuracy obtained in real data.

**Supplementary Fig. 7.**
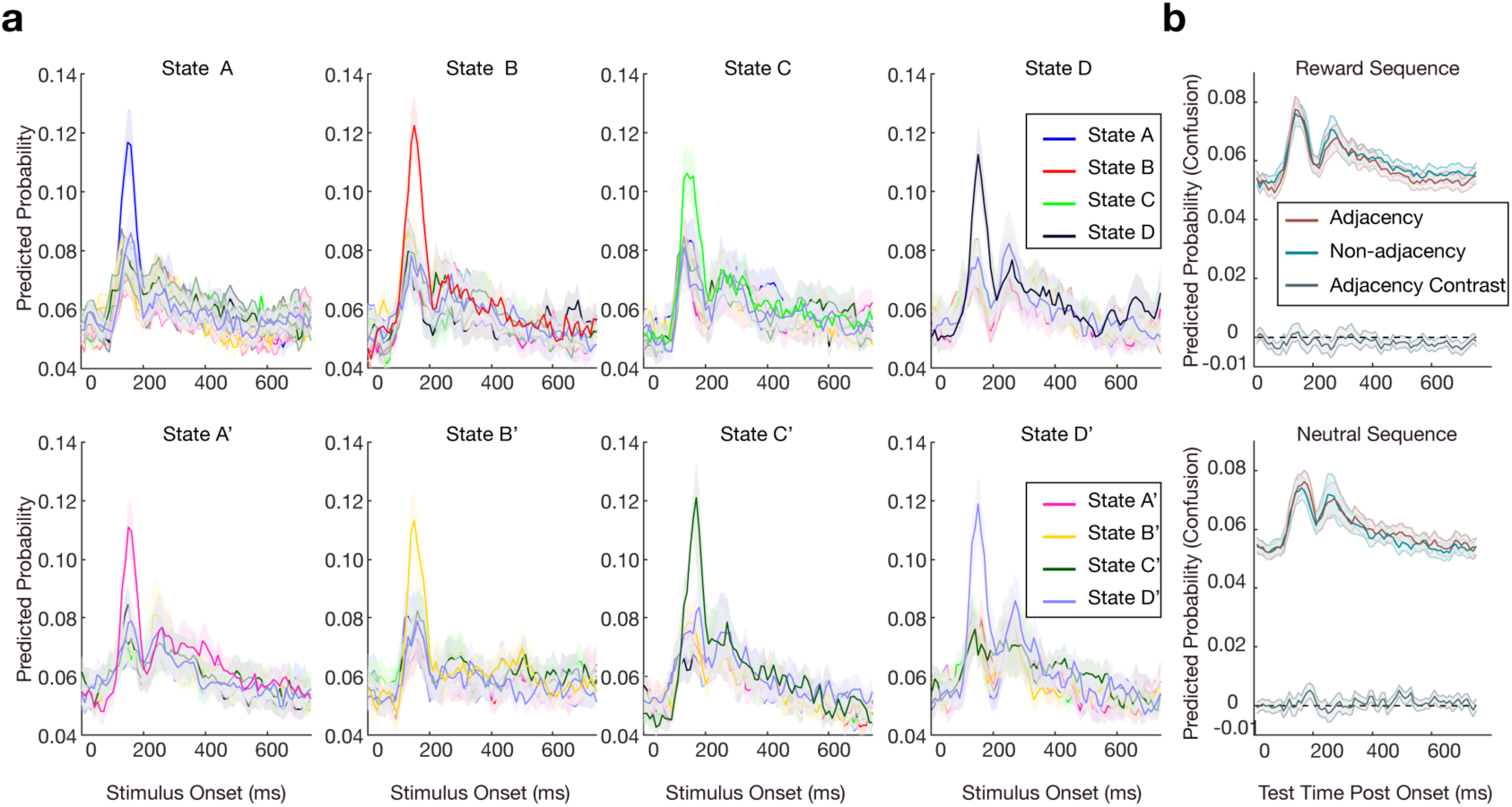
Stability of Neural Decoding and “Confusion” Between Adjacent Task States. **a**, Cross-temporal decoding. Classifiers trained at 180 ms post-stimulus in the functional-localiser phase were tested, across time, on data from the final position-test phase (x-axis). This assesses whether neural patterns learned at the start of the experiment remain discriminable at the end. Line colour denotes the classifier’s target state; each panel plots the true state presented. The sharp, state-specific profiles indicate robust and stable decoding, arguing against substantial representational drift. Shaded bands show 95 % confidence intervals across participants. (For comparison, Supplementary Fig. 1 trains and tests entirely within the localiser phase.) **b,** Classifier “confusion” probabilities for adjacent pairs (e.g., A→B, B→C) versus non-adjacent pairs (e.g., A→C, B→D) in reward (top) and neutral (bottom) sequences. Mixed-effects modelling revealed no adjacency effect and no adjacency × time interaction, indicating that neighbouring states did not become more similar than other state pairs after learning.

**Supplementary Fig. 8.**
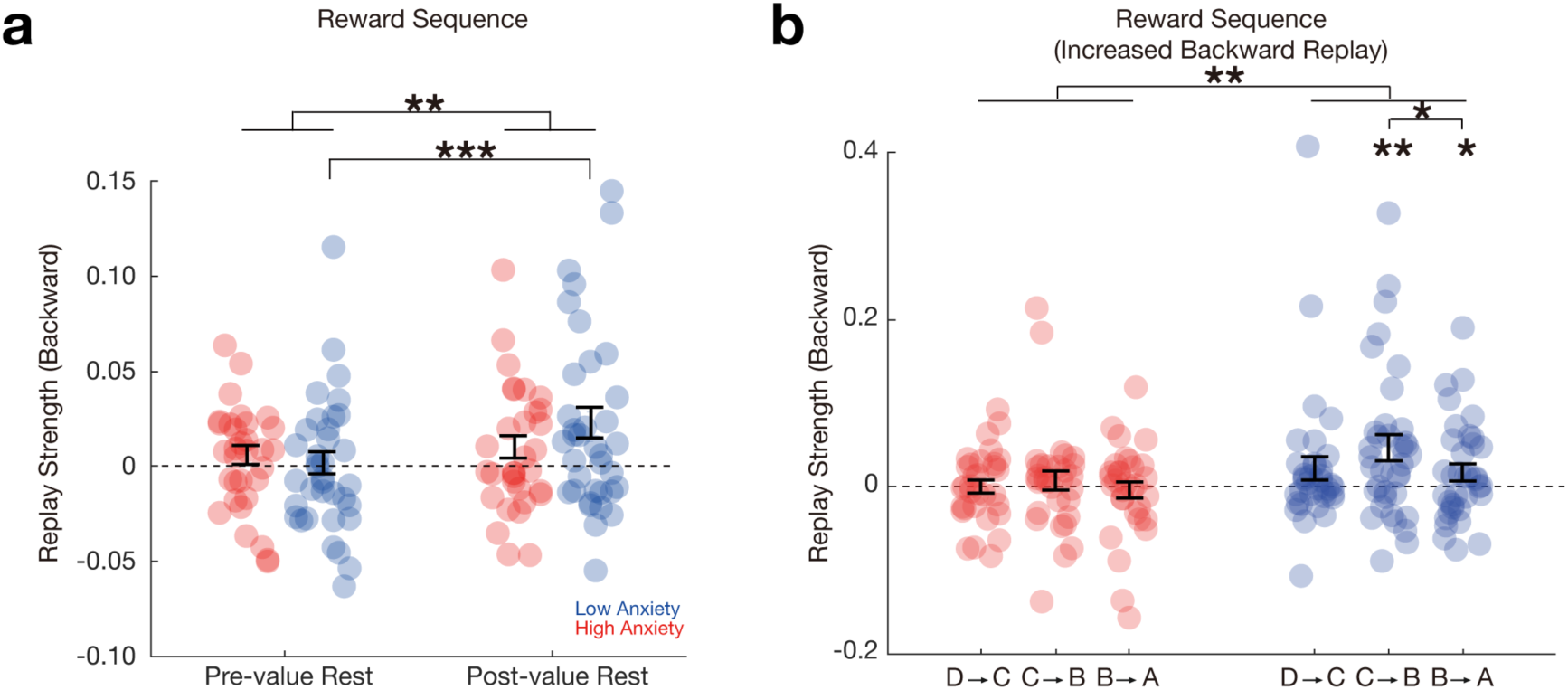
Increase of Replay as a Function of Time and Distance, related to. Figure 4**. a,** There was an interaction effect between group and time for the backward replay, such that individuals with low anxiety exhibited a greater increase of backward replay than those with high anxiety post value learning. **b,** Low anxious subjects (blue) showed significant increase of backward replay for all pairwise associations, with stronger increase for transitions nearer the reward than distal to the reward (paired *t* test between *C* → *B* and *B* → *A*, *t* (35) =1.65, *p* = 0.05), while for subjects with high anxiety, there is no significant replay for any pairwise associations within reward sequence (red). Error bars show SEM; each dot indicates results from one subject. \**p* < 0.05, \*\**p* < 0.01, \*\*\**p* < 0.001.

**Supplementary Fig. 9.**
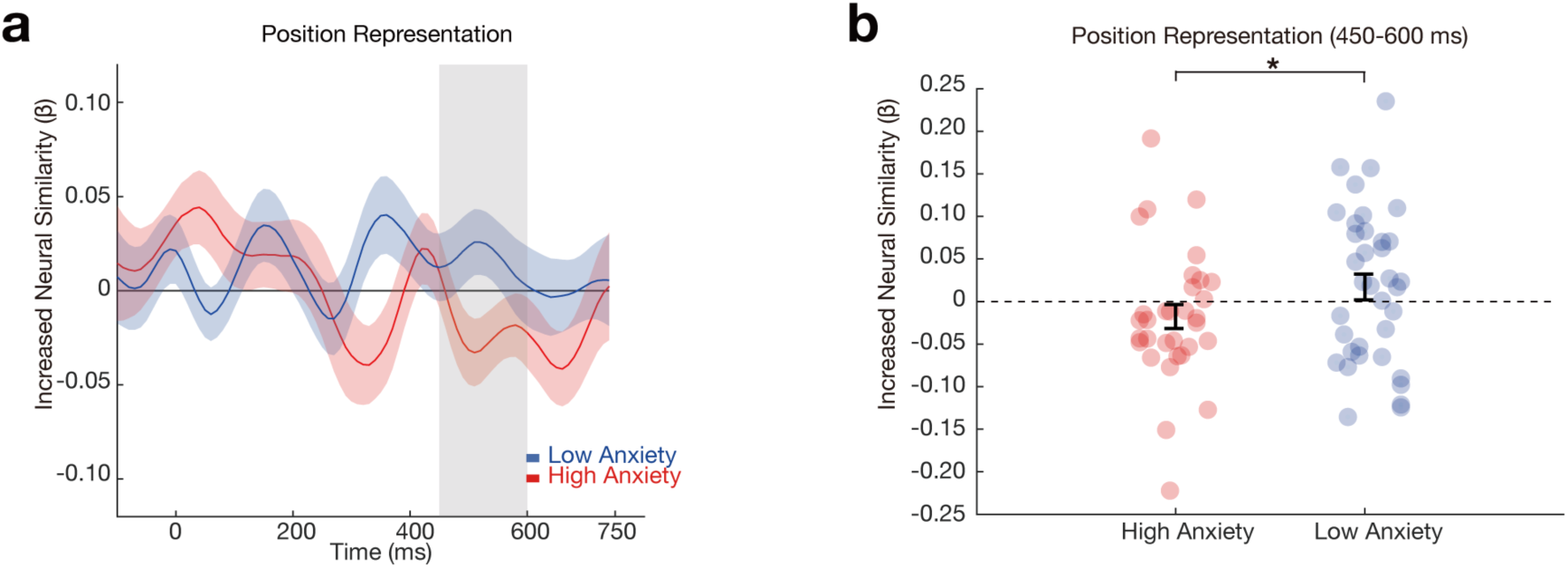
Neural Representation of Position with Anxiety, related to. Figure 5**. a,** The temporal dynamics of the emergent (learning-induced) position representation, separately for low (blue) and high (red) anxious participants. **b,** Based on previous study ^1^, a 450-600ms time window after the stimuli onset (grey shaded area in panel a) was chosen as the region of interest (ROI). The ROI analysis revealed a significant increase of the position representation in low than high anxious subjects (*t*-test, *t* (66) = 1.68, *p* = 0.049). Error bars show SEM; each dot indicates results from one subject. The solid line reflects the best robust linear fit, \**p* < 0.05.

## References

1. Nour, M. M., Liu, Y., Arumuham, A., Kurth-Nelson, Z. & Dolan, R. J. Impaired neural replay of inferred relationships in schizophrenia. Cell 184, 4315–4328.e17 (2021).

2. Sharp, P. B. & Eldar, E. Computational models of anxiety: nascent efforts and future directions. Curr. Dir. Psychol. Sci. 28, 170–176 (2019).

3. Wise, T., Liu, Y., Chowdhury, F. & Dolan, R. J. Model-based aversive learning in humans is supported by preferential task state reactivation. Sci. Adv. 7, eabf9616 (2021).

4. Browning, M., Behrens, T. E., Jocham, G., O’Reilly, J. X. & Bishop, S. J. Anxious individuals have difficulty learning the causal statistics of aversive environments. Nat. Neurosci. 18, 590–596 (2015).

5. Taylor, C. T., Hoffman, Samantha.N. & Khan, Amanda. J. Anhedonia in anxiety disorders. in vol. 58 201–218 (Current topics in behavioral neurosciences, 2022).

6. Dunsmoor, J. E. & Paz, R. Fear generalization and anxiety: behavioral and neural mechanisms. Biol. Psychiatry 78, 336–343 (2015).

7. Grupe, D. W. & Nitschke, J. B. Uncertainty and anticipation in anxiety: an integrated neurobiological and psychological perspective. Nat. Rev. Neurosci. 14, 488– 501 (2013).

8. Paulus, M. P. & Yu, A. J. Emotion and decision-making: affect-driven belief systems in anxiety and depression. Trends Cogn. Sci. 16, 476–483 (2012).

9. Sailer, U. et al. Altered reward processing in the nucleus accumbens and mesial prefrontal cortex of patients with posttraumatic stress disorder. Neuropsychologia 46, 2836–2844 (2008).

10. Sharot, T., Riccardi, A. M., Raio, C. M. & Phelps, E. A. Neural mechanisms mediating optimism bias. Nature 450, 102–105 (2007).

11. Sharot, T. & Garrett, N. Forming beliefs: why valence matters. Trends Cogn. Sci. 20, 25–33 (2016).

12. Liu, Y., Nour, M. M., Schuck, N. W., Behrens, T. E. J. & Dolan, R. J. Decoding cognition from spontaneous neural activity. Nat. Rev. Neurosci. 23, 204–214 (2022).

13. Foster, D. J. Replay comes of age. Annu. Rev. Neurosci. 40, 581–602 (2017).

14. Buzsáki, G. Hippocampal sharp wave-ripple: A cognitive biomarker for episodic memory and planning. Hippocampus 25, 1073–1188 (2015).

15. Lee, A. K. & Wilson, M. A. Memory of sequential experience in the hippocampus during slow wave sleep. Neuron 36, 1183–1194 (2002).

16. Louie, K. & Wilson, M. A. Temporally structured replay of awake hippocampal ensemble activity during rapid eye movement sleep. Neuron 29, 145–156 (2001).

17. Wilson, M. A. & McNaughton, B. L. Reactivation of hippocampal ensemble memories during sleep. Science 265, 676–679 (1994).

18. Liu, Y., Dolan, R. J., Kurth-Nelson, Z. & Behrens, T. E. J. Human replay spontaneously reorganizes experience. Cell 178, 640–652.e14 (2019).

19. Ambrose, R. E., Pfeiffer, B. E. & Foster, D. J. Reverse replay of hippocampal place cells is uniquely modulated by changing reward. Neuron 91, 1124–1136 (2016).

20. Dupret, D., O’Neill, J., Pleydell-Bouverie, B. & Csicsvari, J. The reorganization and reactivation of hippocampal maps predict spatial memory performance. Nat. Neurosci. 13, 995–1002 (2010).

21. Singer, A. C. & Frank, L. M. Rewarded outcomes enhance reactivation of experience in the hippocampus. Neuron 64, 910–921 (2009).

22. Mattar, M. G. & Daw, N. D. Prioritized memory access explains planning and hippocampal replay. Nat. Neurosci. 21, 1609–1617 (2018).

23. Stachenfeld, K. L., Botvinick, M. M. & Gershman, S. J. The hippocampus as a predictive map. Nat. Neurosci. 20, 1643–1653 (2017).

24. Foster, D. J. & Wilson, M. A. Reverse replay of behavioural sequences in hippocampal place cells during the awake state. Nature 440, 680–683 (2006).

25. Barron, H. C. et al. Neuronal computation underlying inferential reasoning in humans and mice. Cell 183, 228–243.e21 (2020).

26. Liu, Y., Mattar, M. G., Behrens, T. E. J., Daw, N. D. & Dolan, R. J. Experience replay is associated with efficient nonlocal learning. Science 372, eabf1357 (2021).

27. Heller, A. S. & Bagot, R. C. Is hippocampal replay a mechanism for anxiety and depression? JAMA Psychiatry 77, 431–432 (2020).

28. Liu, Y. et al. Temporally delayed linear modelling (TDLM) measures replay in both animals and humans. eLife 10, e66917 (2021).

29. Liu, Y., Mattar, M. G., Behrens, T. E. J., Daw, N. D. & Dolan, R. J. Experience replay is associated with efficient nonlocal learning. Science 372, eabf1357 (2021).

30. McFadyen, J., Liu, Y. & Dolan, R. J. Differential replay of reward and punishment paths predicts approach and avoidance. Nat. Neurosci. 26, 627–637 (2023).

31. Wimmer, G. E., Liu, Y., Vehar, N., Behrens, T. E. J. & Dolan, R. J. Episodic memory retrieval success is associated with rapid replay of episode content. Nat. Neurosci. 23, 1025–1033 (2020).

32. Wimmer, G. E., Liu, Y., McNamee, D. C. & Dolan, R. J. Distinct replay signatures for prospective decision-making and memory preservation. Proc. Natl. Acad. Sci. 120, e2205211120 (2023).

33. Schuck, N. W. & Niv, Y. Sequential replay of nonspatial task states in the human hippocampus. Science 364, eaaw5181 (2019).

34. Wittkuhn, L. & Schuck, N. W. Dynamics of fMRI patterns reflect sub-second activation sequences and reveal replay in human visual cortex. Nat. Commun. 12, 1795 (2021).

35. Gagne, C., Zika, O., Dayan, P. & Bishop, S. J. Impaired adaptation of learning to contingency volatility in internalizing psychopathology. eLife 9, e61387 (2020).

36. Weger, M. & Sandi, C. High anxiety trait: A vulnerable phenotype for stress-induced depression. Neurosci. Biobehav. Rev. 87, 27–37 (2018).

37. Zheng, X. et al. A report on state-trait anxiety in Changchun. *Chin*. J. Ment. Health 7, 60–62 (1993).

38. Zheng, X. & Li, Y. State-trait anxiety inventory. Chin. Ment. Health J. 11, 219–220 (1997).

39. Jafarpour, A., Penny, W., Barnes, G., Knight, R. T. & Duzel, E. Working memory replay prioritizes weakly attended events. *eneuro* 4, ENEURO.0171-17.2017 (2017).

40. Ou, J. et al. Replay builds an efficient cognitive map offline to avoid computation online. (2025) doi:10.1101/2025.01.08.632067.

41. Schapiro, A. C., McDevitt, E. A., Rogers, T. T., Mednick, S. C. & Norman, K. A. Human hippocampal replay during rest prioritizes weakly learned information and predicts memory performance. Nat. Commun. 9, 3920 (2018).

42. Wimmer, G. E. & Shohamy, D. Preference by association: how memory mechanisms in the hippocampus bias decisions. Science 338, 270–273 (2012).

43. Zung, W. W. K. A rating instrument for anxiety disorders. Psychosomatics 12, 371–379 (1971).

44. Meyer, T. J., Miller, M. L., Metzger, R. L. & Borkovec, T. D. Development and validation of the penn state worry questionnaire. Behav. Res. Ther. 28, 487–495 (1990).

45. Zhong, J., Wang, C., Li, J. & Liu, J. Penn State Worry Questionnaire: structure and psychometric properties of the Chinese version. J. Zhejiang Univ. Sci. B 10, 211–218 (2009).

46. Carleton, R. N., Norton, M. A. P. J. & Asmundson, G. J. G. Fearing the unknown: A short version of the Intolerance of Uncertainty Scale. J. Anxiety Disord. 21, 105–117 (2007).

47. Zhang Y., et al. Reliability and validity of the intolerance of uncertainty scale-short form in university studen. Chin. J. Chinical Psychol. 25, 280–288 (2017).

48. Zung, W. W. K. A Self-Rating Depression Scale. Arch. Gen. Psychiatry 12, 63 (1965).

49. Knowles, K. A. & Olatunji, B. O. Specificity of trait anxiety in anxiety and depression: Meta-analysis of the State-Trait Anxiety Inventory. Clin. Psychol. Rev. 82, 101928 (2020).

50. Sandi, C. & Richter-Levin, G. From high anxiety trait to depression: a neurocognitive hypothesis. Trends Neurosci. 32, 312–320 (2009).

51. Shadli, S. M. et al. Right frontal anxiolytic-sensitive EEG ‘theta’ rhythm in the stop-signal task is a theory-based anxiety disorder biomarker. Sci. Rep. 11, 19746 (2021).

52. Poldrack, R. A., Huckins, G. & Varoquaux, G. Establishment of best practices for evidence for prediction: a review. JAMA Psychiatry 77, 534–540 (2020).

53. Bates, D., Kliegl, R., Vasishth, S. & Baayen, H. Parsimonious mixed models. Preprint at arXiv:1506.04967 (2015).

54. Bates, D., Mächler, M., Bolker, B. & Walker, S. Fitting Linear Mixed-Effects Models Using lme4. J. Stat. Softw. 67, (2015).

55. Gelman, A. & Brown, N. J. L. How statistical challenges and misreadings of the literature combine to produce unreplicable science: An example from psychology. Adv. Methods Pract. Psychol. Sci. 7, 25152459241276398 (2024).

56. Usui, Y. et al. Effects of inappropriate nurturing experiences, depressive Rumination, and trait anxiety on depressive symptoms. Neuropsychiatr. Dis. Treat. Volume 20, 1571–1581 (2024).

57. Wang, T. et al. Relations between trait anxiety and depression: A mediated moderation model. J. Affect. Disord. 244, 217–222 (2019).

58. Luyckx, F., Nili, H., Spitzer, B. & Summerfield, C. Neural structure mapping in human probabilistic reward learning. eLife 7, e42816 (2019).

59. Spielberger, C. D., Gorsuch, R., Lushene, R., Vagg, P. & Jacobs, G. Manual for the State-Trait Anxiety Inventory. vol. IV (Consulting Psychologists Press, 1983).

60. Spielberger, C. D., Gonzalez-Reigosa, F. & Martinez-Urrutia, A. Development of the spanish edition of the state-trait anxiety inventory. Interam. J. Psychol. 3–4 (1971).

61. Pike, A. C. & Robinson, O. J. Reinforcement learning in patients with mood and anxiety disorders vs control individuals: a systematic review and meta-analysis. JAMA Psychiatry 79, 313–322 (2022).

62. Eckstrand, K. L. et al. Trauma-associated anterior cingulate connectivity during reward learning predicts affective and anxiety states in young adults. Psychol. Med. 49, 1831–1840 (2019).

63. Liu, Q. et al. Neural function underlying reward expectancy and attainment in adolescents with diverse psychiatric symptoms. NeuroImage Clin. 36, 103258 (2022).

64. Auerbach, R. P. et al. Reward-related neural circuitry in depressed and anxious adolescents: a human connectome project. J. Am. Acad. Child Adolesc. Psychiatry 61, 308–320 (2022).

65. Anderson, Z. et al. Association between reward-related functional connectivity and tri-level mood and anxiety symptoms. NeuroImage Clin. 37, 103335 (2023).

66. Etkin, A. & Fonzo, G. A. Learning in generalized anxiety disorder benefits from neither the carrot nor the stick. Am. J. Psychiatry 174, 87–88 (2017).

67. Anacker, C. & Hen, R. Adult hippocampal neurogenesis and cognitive flexibility — linking memory and mood. Nat. Rev. Neurosci. 18, 335–346 (2017).

68. Fung, B. J., Qi, S., Hassabis, D., Daw, N. & Mobbs, D. Slow escape decisions are swayed by trait anxiety. *Nat*. Hum. Behav. 3, 702–708 (2019).

69. Gorka, A. X., Hanson, J. L., Radtke, S. R. & Hariri, A. R. Reduced hippocampal and medial prefrontal gray matter mediate the association between reported childhood maltreatment and trait anxiety in adulthood and predict sensitivity to future life stress. Biol. Mood Anxiety Disord. 4, 12 (2014).

70. Nemat Tavousi, M. & Seyf Hashemi, N. The relationship between perfectionism and depression and social anxiety in social media users: Emphasizing the mediating role of rumination. J. Assess. Res. Appl. Couns. 6, 152–160 (2024).

71. Nolen-Hoeksema, S. Responses to depression and their effects on the duration of depressive episodes. J. Abnorm. Psychol. 100, 569–582 (1991).

72. Nolen-Hoeksema, S. The role of rumination in depressive disorders and mixed anxiety/depressive symptoms. J. Abnorm. Psychol. 109, 504–511 (2000).

73. LeDoux, J. E. Coming to terms with fear. Proc. Natl. Acad. Sci. 111, 2871–2878 (2014).

74. LeDoux, J. E. As soon as there was life, there was danger: the deep history of survival behaviours and the shallower history of consciousness. Philos. Trans. R. Soc. B Biol. Sci. 377, 20210292 (2022).

75. Michely, J., Rigoli, F., Rutledge, R. B., Hauser, T. U. & Dolan, R. J. Distinct processing of aversive experience in amygdala subregions. Biol. Psychiatry Cogn. Neurosci. Neuroimaging 5, 291–300 (2020).

76. Voegler, R., Peterburs, J., Bellebaum, C. & Straube, T. Modulation of feedback processing by social context in social anxiety disorder (SAD)-an event-related potentials (ERPs) study. Sci. Rep. 9, 4795 (2019).

77. Yamamori, Y., Robinson, O. J. & Roiser, J. P. Approach-avoidance reinforcement learning as a translational and computational model of anxiety-related avoidance. eLife 12, RP87720 (2023).

78. Sharp, P. B., Dolan, R. J. & Eldar, E. Disrupted state transition learning as a computational marker of compulsivity. Psychol. Med. 53, 2095–2105 (2023).

79. Sharp, P. B. & Eldar, E. Computational models of anxiety: Nascent efforts and future directions. Curr. Dir. Psychol. Sci. 28, 170–176 (2019).

80. Nour, M. M., Liu, Y., Higgins, C., Woolrich, M. W. & Dolan, R. J. Reduced coupling between offline neural replay events and default mode network activation in schizophrenia. Brain Commun. 5, fcad056 (2023).

81. Nour, M. M., McNamee, D. C., Liu, Y. & Dolan, R. J. Trajectories through semantic spaces in schizophrenia and the relationship to ripple bursts. Proc. Natl. Acad. Sci. 120, e2305290120 (2023).

82. Sommer, V. R. et al. Neural pattern similarity differentially relates to memory performance in younger and older adults. J. Neurosci. 39, 8089–8099 (2019).

83. Sommer, V. R., Mount, L., Weigelt, S., Werkle-Bergner, M. & Sander, M. C. Spectral pattern similarity analysis: Tutorial and application in developmental cognitive neuroscience. Dev. Cogn. Neurosci. 54, 101071 (2022).

84. Gurariy, G., Mruczek, R. E. B., Snow, J. C. & Caplovitz, G. P. Using high-density electroencephalography to explore spatiotemporal representations of object categories in visual cortex. J. Cogn. Neurosci. 34, 967–987 (2022).

85. Hsu, S.-H., Lin, Y., Onton, J., Jung, T.-P. & Makeig, S. Unsupervised learning of brain state dynamics during emotion imagination using high-density EEG. NeuroImage 249, 118873 (2022).

86. Huang, Q. et al. Replay-triggered brain-wide activation in humans. Nat. Commun. 15, 7185 (2024).

87. Zika, O., Wiech, K., Reinecke, A., Browning, M. & Schuck, N. W. Trait anxiety is associated with hidden state inference during aversive reversal learning. Nat. Commun. 14, 4203 (2023).

88. Moran, T. P. Anxiety and working memory capacity: A meta-analysis and narrative review. Psychol. Bull. 142, 831–864 (2016).

89. Fisher, P. L. & Durham, R. C. Recovery rates in generalized anxiety disorder following psychological therapy: an analysis of clinically significant change in the STAI-T across outcome studies since 1990. Psychol. Med. 29, 1425–1434 (1999).

90. Hein, T. P. et al. Anterior cingulate and medial prefrontal cortex oscillations underlie learning alterations in trait anxiety in humans. *Commun*. Biol. 6, 271 (2023).

91. Eysenck, M. W., Moser, J. S., Derakshan, N., Hepsomali, P. & Allen, P. A neurocognitive account of attentional control theory: how does trait anxiety affect the brain’s attentional networks? Cogn. Emot. 37, 220–237 (2023).

92. Kurth-Nelson, Z., Barnes, G., Sejdinovic, D., Dolan, R. & Dayan, P. Temporal structure in associative retrieval. eLife 4, e04919 (2015).

93. Kurth-Nelson, Z., Economides, M., Dolan, R. J. & Dayan, P. Fast sequences of non-spatial state representations in humans. Neuron 91, 194–204 (2016).

